# Lytic infection with murine gammaherpesvirus 68 activates host and viral RNA polymerase III promoters and enhances non-coding RNA expression

**DOI:** 10.1101/795278

**Authors:** Ashley N. Knox, Alice Mueller, Eva M. Medina, Eric T. Clambey, Linda F. van Dyk

**Author notes:** Address correspondence to Linda van Dyk.

## Abstract

RNA polymerase III (pol III) transcribes multiple non-coding (nc) RNAs that are essential for cellular function. Pol III-dependent transcription is also engaged during certain viral infections, including the gammaherpesviruses (γHVs), where pol III-dependent viral ncRNAs promote pathogenesis. Additionally, several host ncRNAs are upregulated during γHV infection and play integral roles in pathogenesis by facilitating viral establishment and gene expression. Here, we sought to investigate how pol III promoters and transcripts are regulated during gammaherpesvirus infection using the murine gammaherpesvirus 68 (γHV68) system. To compare the transcription of host and viral pol III-dependent ncRNAs, we analyzed a series of pol III promoters for host and viral ncRNAs using a luciferase reporter optimized to measure pol III activity. We measured promoter activity from the reporter gene at the translation level via luciferase activity and at the transcription level via RT-qPCR. We further measured endogenous ncRNA expression at single cell-resolution by flow cytometry. These studies demonstrated that lytic infection with γHV68 increased the transcription from multiple host and viral pol III promoters, and further identified the ability of accessory sequences to influence both baseline and inducible promoter activity after infection. RNA flow cytometry revealed the induction of endogenous pol III-derived ncRNAs that tightly correlated with viral gene expression. These studies highlight how lytic gammaherpesvirus infection alters the transcriptional landscape of host cells to increase pol III-derived RNAs, a process that may further modify cellular function and enhance viral gene expression and pathogenesis.

**IMPORTANCE:** Gammaherpesviruses are a prime example of how viruses can alter the host transcriptional landscape to establish infection. Despite major insights into how these viruses modify RNA polymerase II-dependent generation of messenger RNAs, how these viruses influence the activity of host RNA polymerase III remains much less clear. Small non-coding RNAs produced by RNA polymerase III are increasingly recognized to play critical regulatory roles in cell biology and virus infection. Studies of RNA polymerase III dependent transcription are complicated by multiple promoter types and diverse RNAs with variable stability and processing requirements. Here, we characterized a reporter system to directly study RNA polymerase III-dependent responses during gammaherpesvirus infection and utilized single-cell flow cytometry-based methods to reveal that gammaherpesvirus lytic replication broadly induces pol III activity to enhance host and viral non-coding RNA expression within the infected cell.

## INTRODUCTION

Gammaherpesviruses (γHVs) are large, dsDNA viruses that establish a life-long infection in their hosts, with long-term latency in lymphocytes (1, 2). The γHVs include the human pathogens, Epstein-Barr virus (EBV) and Kaposi’s sarcoma-associated herpesvirus (KSHV or HHV-8), and murine gammaherpesvirus 68 (γHV68 or MHV-68; ICTV nomenclature *Murid herpesvirus 4*, MuHV-4) (3). These viruses establish a primary lytic infection in their host that is followed by a prolonged quiescent infection termed latency. Latency is maintained in healthy individuals by a homeostatic relationship between the virus and the host immune response; if this balance is disrupted (e.g. by immunosuppression), γHVs can reactivate from latency and actively replicate. Disruption between the balance of γHV infection and host immune control is associated with γHV multiple pathologies, including a range of malignancies (4).

The γHVs contain several types of non-coding (nc) RNAs, including nuclear ncRNAs and functional miRNAs; these diverse RNAs include ncRNAs transcribed by RNA polymerase II (e.g. the KSHV PAN RNA and the KSHV and EBV miRNAs) or by RNA pol III (e.g. the EBV-encoded small RNAs (EBERs) and the γHV68 tRNA-miRNA-encoded RNAs (TMERs)) (5-13). Viral ncRNAs are considered to have important host-modulatory functions, interacting with host proteins and regulating host and viral gene expression. For example, the EBV EBERs are expressed during latency, and were discovered through their interaction with the host lupus-associated antigen (La) protein, which putatively mediates EBER interaction with TLR3 (14-17).The EBERs have further been shown to interact with several host proteins including ribosomal protein L22, protein kinase R (PKR), and retinoic-acid inducible gene I (RIG-I) (18). These interactions can trigger sustained host innate immune responses that are implicated in the development of EBV-associated malignancies (16, 19-21). γHV68, a highly tractable small animal model of γHV infection, also encodes several pol III-transcribed ncRNAs known as the tRNA-miRNA-encoded RNAs (TMERs) (22, 23). The TMERs are dispensable for lytic replication and establishment of latency; however, these transcripts are required for pathogenesis during acute infection of an immunocompromised host (7, 24, 25). The TMERs contain bi-functional elements with a tRNA-like structure at the 5’ end and hairpins that are processed into biologically-active miRNAs (7), capable of targeting a number of RNAs for post-transcriptional regulation (26). Our lab has previously shown that the tRNA-like structure is sufficient to rescue pathogenesis of a TMER deficient viral recombinant, suggesting that like the EBERs, the TMERs may contribute to pathogenesis through their interactions with host proteins (25). Though TMER-host protein interactions have yet to be fully explored, it is notable that several characteristics of the EBERs, such as a 5’-triphosphate and 3’-polyU, are imparted by RNA polymerase III (pol III) transcription (27).These motifs can be recognized by host RNA-binding proteins, such as RIG-I or La, to trigger an innate immune response (16, 20, 27, 28).

Pol III is often considered to perform “house-keeping” functions, as it transcribes host genes required for cell growth and maintenance (e.g. U6 snRNA, tRNAs, and 5S rRNA) (29). Despite this, it is clear that the γHVs can usurp pol III-dependent transcription mechanisms for their own purposes. Latent EBV infection has been shown to upregulate components of pol III and ultimately increase the expression of host pol III transcripts – particularly vault RNAs – that allow increased establishment of viral infection and gene expression (30-32). Similarly, γHV68 infection drives upregulation of host pol III-dependent short-interspersed nuclear element (SINE) RNAs, which in turn, mediate increased viral gene expression (33-35). Additionally, our lab has reported that reactivation of a latently-infected γHV68 cell line results in a rare subset of the population that demonstrates increased viral transcription and translation, including increased expression of TMERs (36). Notably, dysregulation of pol III is a common feature of many cancer cells, implicating γHV infection-driven alteration of pol III activity as one potential contributor to γHV-associated malignancies (37). Therefore, understanding how γHV infection alters pol III activity is integral to elucidating mechanisms of γHV pathogenesis.

The analysis of pol III activity during γHV infection has been complicated by the nature of pol III-derived ncRNAs. These transcripts are often short and structured, creating complications in probe specificity to quantitatively analyze promoter activity/gene expression by conventional means (e.g. northern blot, RT-qPCR). Probe specificity is also challenging for classes of RNAs with highly conserved promoter features, such as the γHV68 TMERs or the human tRNAs. However, promoter analysis of the TMERs could reveal rules of transcription that apply to other conserved and potentially co-regulated ncRNAs, such as the human tRNAs. Additionally, many pol III-derived ncRNAs may be scarce or abundant so that changes in expression can be obscured. Therefore, highly sensitive readouts, such as the high dynamic range of luciferase assays or quantitative assays with single-cell resolution, offer potential improvement in measuring pol III-derived ncRNA expression. Single cell RNA flow cytometry allows RNA detection with high specificity without the need for unique primers and probe, and has the additional benefit of measuring RNA levels in individual cells to reveal fine fluctuations that may be obscured in bulk analyses.

The ideal comparison of promoters allows variation only in the promoter elements coupled to a common reporter gene and sensitive detection. Traditional analysis of RNA pol II promoter activity has benefitted significantly from the use of luciferase reporter systems, which provide the advantages of a readout that is high throughput, has a wide dynamic range for maximal quantitation, and is standardized across varied promoters and cellular conditions. RNA pol III does not produce coding RNAs in normal biology; however, several studies have reported the use of luciferase reporters to measure ncRNA derived from pol III or pol I (38, 39). Based on these studies, we developed a panel of luciferase reporters driven by pol III promoters to determine the efficacy of a reporter gene approach in analyzing ncRNA promoter activity during viral infection. As with analysis of RNA pol II reporters, caveats to the enzymatic readout of this system are that it is several steps downstream of RNA transcription, the efficacy of RNA translation may differ among specific RNAs, and infection may alter translation in a number of ways yet to be described. However, use of the facile enzymatic readout plus RT-qPCR quantitation of the reporter RNA allows us to directly compare these measures for highly sensitive quantitative analysis. As a complementary approach, we quantified ncRNAs expression at the single-cell level in the presence or absence of virus infection.

Due to the importance of γHV ncRNAs during infection and the unique transcriptional regulation afforded by RNA pol III, the overall objective of this study was to compare different RNA pol III promoters and their activity during virus infection using three different methods for sensitive and quantitative analysis. We found that γHV68 infection upregulates the activity of multiple viral and host pol III promoters, a process further associated with the induction of pol III-dependent targets. These studies indicate that lytic γHV infection can broadly enhance RNA pol III promoter activity to modify the ncRNA landscape of infected cells.

## RESULTS

RNA polymerase III can transcribe RNA from a variety of gene-internal (type 1 and 2) and gene-external (type 3) promoters (Fig 1A). These promoters contain distinct motifs that determine which transcription factors bind to the promoter to recruit pol III (40). To understand how γHV lytic replication influenced RNA pol III promoter activity while limiting the confounding factors of the individual RNA primary transcripts and modified or processed products, we sought to make use of a luciferase assay previously used to study pol III promoter activity (41) to study a series of viral and host pol III promoters. We selected the reporter plasmid pNL1.1 (Promega), because NanoLuc luciferase creates a brighter signal, and the protein is smaller than other luciferase proteins (NanoLuc 19.1 kDa and 171 nucleotides; Renilla 36.0 kDa and 312 nucleotides; Firefly 60.6 kDa and 550 nucleotides), which is consistent with pol III processivity of small ncRNAs. Our analysis of the pNL1.1 sequence revealed a pol III termination signal within the luciferase coding gene (TTTT). Therefore, to examine the activity of pol III promoters without the potential for early termination, we introduced silent mutations into the NanoLuc reporter construct to remove the termination signal. This altered vector was named “pNLP3” to reflect that it is a *<u>NanoLuc</u>* reporter optimized for *<u>p</u>*ol *<u>III</u>* (Fig 1B). The human U6 promoter was cloned into both the parental pNL1.1 and the modified pNLP3 to compare the effects of removing the pol III termination signal, with promoter activity measured 24 hours post-infection. We found that removal of the termination signal increased the luciferase output, indicating that there was more read-through of the full NanoLuc gene from pNLP3 (Fig 1B, left). Furthermore, RT-PCR analysis of the NanoLuc transcript transcribed from the U6 promoter in either the pNL1.1 or pNLP3 vector revealed more full-length NanoLuc transcript from the pNLP3 vector (Fig 1B, right). This indicates that the pNLP3 vector allows for optimal pol III transcription of the reporter gene. We therefore used the pNLP3 vector as the backbone for analysis of all other pol III promoters included in this study.

**Figure 1.**
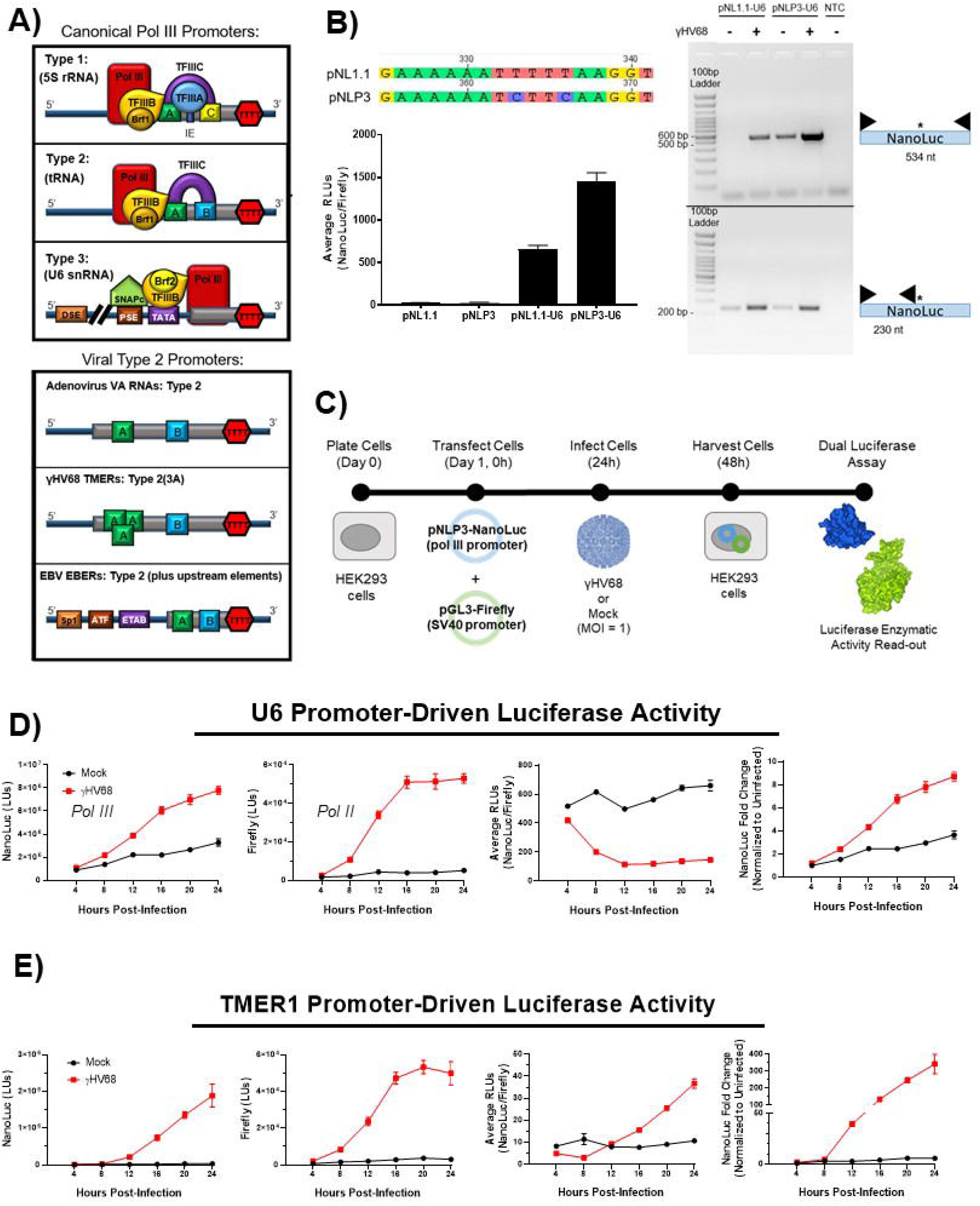
RNA Polymerase III promoters contain distinct motifs and respond to viral infection. **(A)** Pol III transcribes RNA from gene-internal (type 1 and type 2) or gene-external (type 3) promoters. Each promoter type contains distinct motifs, which are in turn bound by specific transcription factors (TF) that recruit pol III to the promoter. Viral promoters can have canonical type 2 promoters, such as adenovirus; however, the γHV68 TMERs contain a triplicated, overlapping A box motif, and the EBV EBER promoters include upstream elements. Gray boxes indicate the gene, while the “TTTT” marks the end of the gene where pol III transcription is terminated. Diagrams are not to scale. A = A box, B = B box, C = C box, IE = intermediate element, “TTTT” = pol III termination signal, TF = transcription factor, TATA = TATA box, PSE = proximal sequence element, DSE = distal sequence elements, ETAB = EBER TATA-like box, ATF = activating transcription factor binding sequence, Sp1 = Sp1-like binding sequence. **(B)** Optimization of the NanoLuc reporter vector. Mutations were introduced into the pNL1.1 vector to remove the pol III termination signal (TTTT) from the NanoLuc coding region, resulting in the pNLP3 vector. The human U6 promoter was cloned into each of these reporters, and dual luciferase assays were performed to compare luciferase output (left panel). Data shown is representative of two independent experiments with biological triplicates. Additionally, RNA was isolated from cells transfected with these two constructs. Cells were mock or WT γHV68-infected for 16 h, then cellular RNA was used as a template for primers targeting the entire NanoLuc gene (top gel, 534 nt), or targeting just the NanoLuc sequence upstream of the termination sequence (bottom gel, 234 nt) (right panel, with primers as indicated). Data shown is from one experiment with biological triplicates. NTC = Non-template control, black triangles = primers, * = location of termination sequence in pNL1.1. **(C)** Experimental design. HEK 293 cells are transfected with a control Firefly reporter (pGL3) and a pNLP3 reporter expressed from a pol III promoter of interest. 24 h post-transfection, cells are infected with wild-type (WT) γHV68 at an MOI of 1, or mock treated. Cell lysates are collected 24 h post-transfection for the dual luciferase assay. Promoter activity was measured for the **(D)** U6 and **(E)** TMER1 promoters using a dual luciferase assay. Cells were harvested every 4 hours post-infection up to 24 h. Each time-point was repeated for biological triplicates. Raw values for the Firefly and NanoLuc activity for each promoter are shown as luminescence units (LUs). Promoter activity was analyzed as average relative luminescence units (RLUs = NanoLuc LUs / Firefly LUs), and as the fold change in NanoLuc luminescence units compared to the 4 h uninfected samples (NanoLuc fold change = Infected Sample NanoLuc LU / Mock Sample NanoLuc LU). Error bars = SEM.

With an optimized pol III reporter construct, we assessed how different pol III promoters respond to γHV68 infection over time. We first compared the activity of the human U6 (type 3) and γHV68 TMER1 (type 2) promoters. HEK 293 cells were transfected with pNLP3 vectors containing either the U6 or TMER1 promoter, co-transfected with an SV40 (pol II promoter)-driven Firefly luciferase vector, then infected with γHV68 (Fig 1C). Cell lysates were collected every 4 hours for 24 hours post-infection to quantify promoter activity as defined by NanoLuc luciferase activity. This analysis revealed that γHV68 infection resulted in a time-dependent increase in NanoLuc activity for both the U6 and TMER1 promoters (left panel, Fig 1D-1E) relative to mock-infected samples. γHV68 infection also resulted in a time-dependent increase in expression of the control, Firefly luciferase reporter (second panel from left, Fig 1D-1E). The results for dual luciferase assays are typically reported as relative luminescence units (RLUs), where the reporter luminescence units (LUs) are normalized to the luminescence units of the control luciferase (i.e., NanoLuc LUs / Firefly LUs). However, since γHV68 infection simultaneously increased luminescence from both the NanoLuc reporter, and from the control Firefly reporter, this normalization implied decreased relative U6 promoter activity with infection when we actually observe an increase in NanoLuc activity (Fig 1D). Clearly, the numerous changes incurred in cells during viral infection limits our ability to standardize pol III promoter activity relative to a pol II promoter control (i.e. SV40 promoter); therefore, all subsequent analyses report promoter activity as a fold change in NanoLuc luminescence comparing mock and γHV68 infected samples. This allows us to directly compare the effect of infection on the reporter in related samples. We found that the U6 promoter drives high basal luciferase activity under mock conditions (left panel, Fig 1D), with a further increase in raw and normalized U6-expressed NanoLuc LUs throughout infection (Fig 1D). In contrast, the TMER1 promoter was characterized by extremely low basal luciferase activity (left panel, Fig 1E) in mock conditions; however, this promoter was strongly induced by infection (Fig 1E). These data suggest that the luciferase assay can be used for analysis of pol III promoters and show that γHV68 lytic infection increases the activity of multiple pol III promoter types, with a more robust induction of the type 2 promoter of TMER1 compared to the U6 promoter.

We further analyzed the activity of several other promoter types to assess how they are impacted during γHV68 infection. Experiments indicated that γHV68 infection induced activity from multiple pol III promoters, including the human U6 and tRNA-Tyr promoters, the EBER1 and EBER2 promoters, and TMER1, 4, and 5 promoters (Fig 2). Though the vaultRNA1-1 and adenovirus VA1 promoters were cloned into the reporter, there was no detectable activity from these constructs (unpublished data). While viral infection induced luciferase activity from all of the examined promoters, the TMER promoters consistently showed the greatest induction during infection.

**Figure 2.**
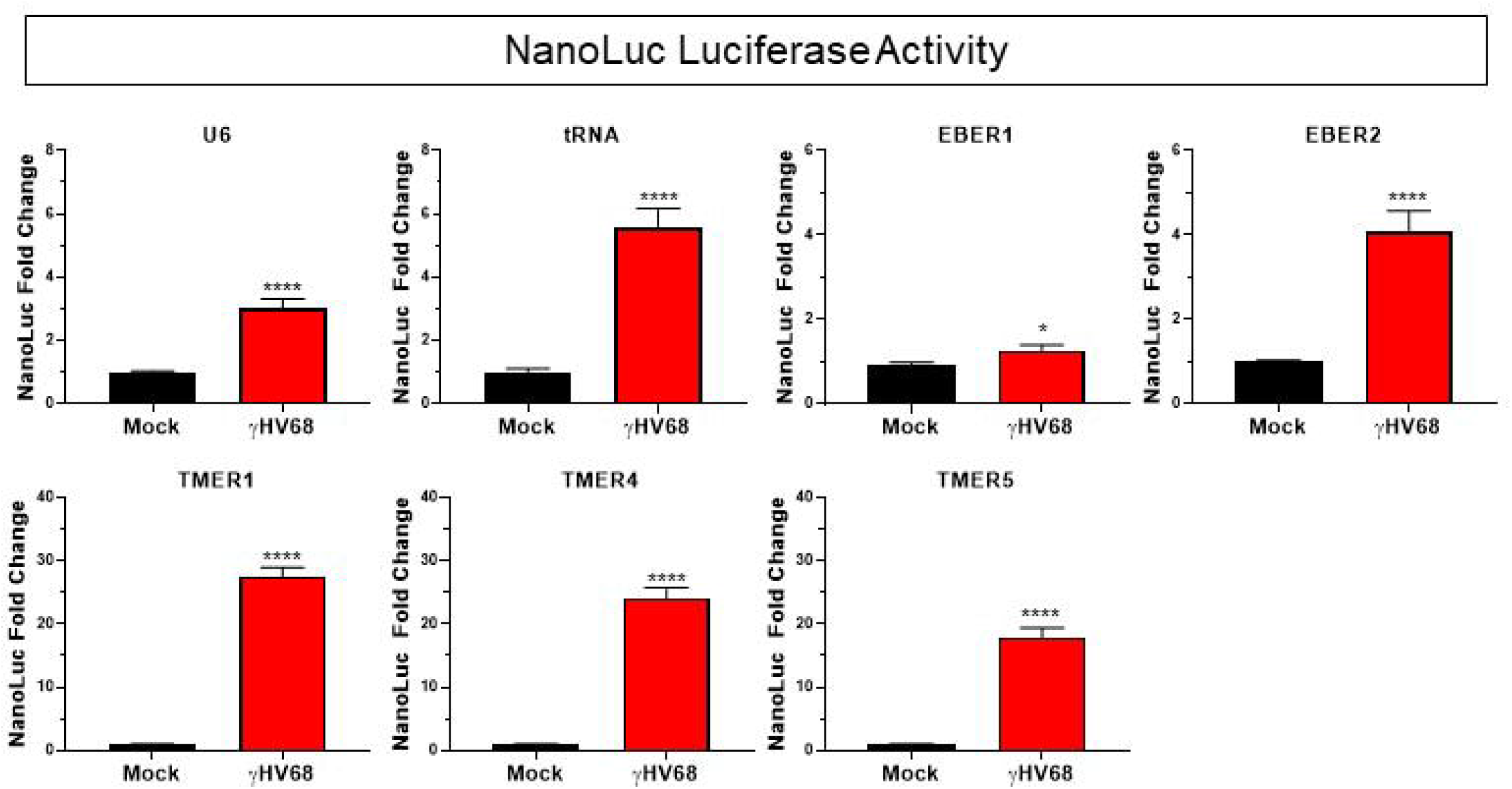
γHV68 infection induces activity of multiple pol III promoter types. Promoter activity was measured for the U6 (n = 8), tRNA (n = 3), EBER1 (n = 3), EBER2 (n = 4), TMER1 (n = 18), TMER4 (n = 4), and TMER5 (n = 6) promoters during infection. Luciferase assays were performed as previously described, with cell lysates collected 24 h post-infection. Each independent experiment (n) contained biological triplicates or duplicates. All promoter activity changes are analyzed as the fold change in NanoLuc activity normalized to uninfected samples (Infected NanoLuc LU / Mock NanoLuc LU). Error bars = SEM. Significant differences analyzed by t-test and indicated as asterisks. P-values are indicated as follows: * = P ≤ 0.05, **** = P ≤ 0.0001.

Because the normal role of pol III is in transcription of non-coding RNAs, we wanted to directly measure the impact of a pol III specific inhibitor (CAS 577784-91-9) on luciferase activity from the TMER1 promoter compared to pol II promoters (TK-NanoLuc and SV40-Firefly). We treated cells with 40 μM of CAS 577784-91-9, a drug concentration previously reported in investigation of γHV68 induction of SINE RNAs (33), prior to transfection and infection. We found that inhibition of pol III with this drug concentration significantly reduced the induction of luciferase activity from the TMER1 promoter without toxicity (unpublished data), and reduced expression of endogenous pol III-transcribed genes during infection (Fig 3A, human pre-tRNA-Tyr-GTA-1-1 and γHV68 TMER1). Pol III inhibition did not affect an endogenous pol II-transcribed gene (Fig 3A, NFAT5) and had no consistent effect on the luciferase activity from pol II promoters SV40 and TK (Fig 3B). In contrast, pol III inhibition led to significant decreases in luciferase activity from six of the seven tested pol III promoters (Fig 3C), with the exception being the U6 promoter (the only type 1 promoter) which exhibited no decrease in activity. The apparent resistance of the U6 promoter to inhibition could either be due to the relatively high activity of the U6 promoter, or to alternate mechanisms of transcription at the U6 promoter (e.g. pol II recruitment to the promoter (41)) invoked at these inhibitor concentrations. Though these data show pol III activity was not completely inhibited, we expect that full inhibition of pol III would result in cell death, and therefore, partial pol III inhibition is ideal for measuring the impact of pol III in this system. These studies indicate that inhibiting pol III activity consistently impaired induction of luciferase from multiple host and viral pol III promoters corresponding to the type 2 pol III promoter subtype.

**Figure 3.**
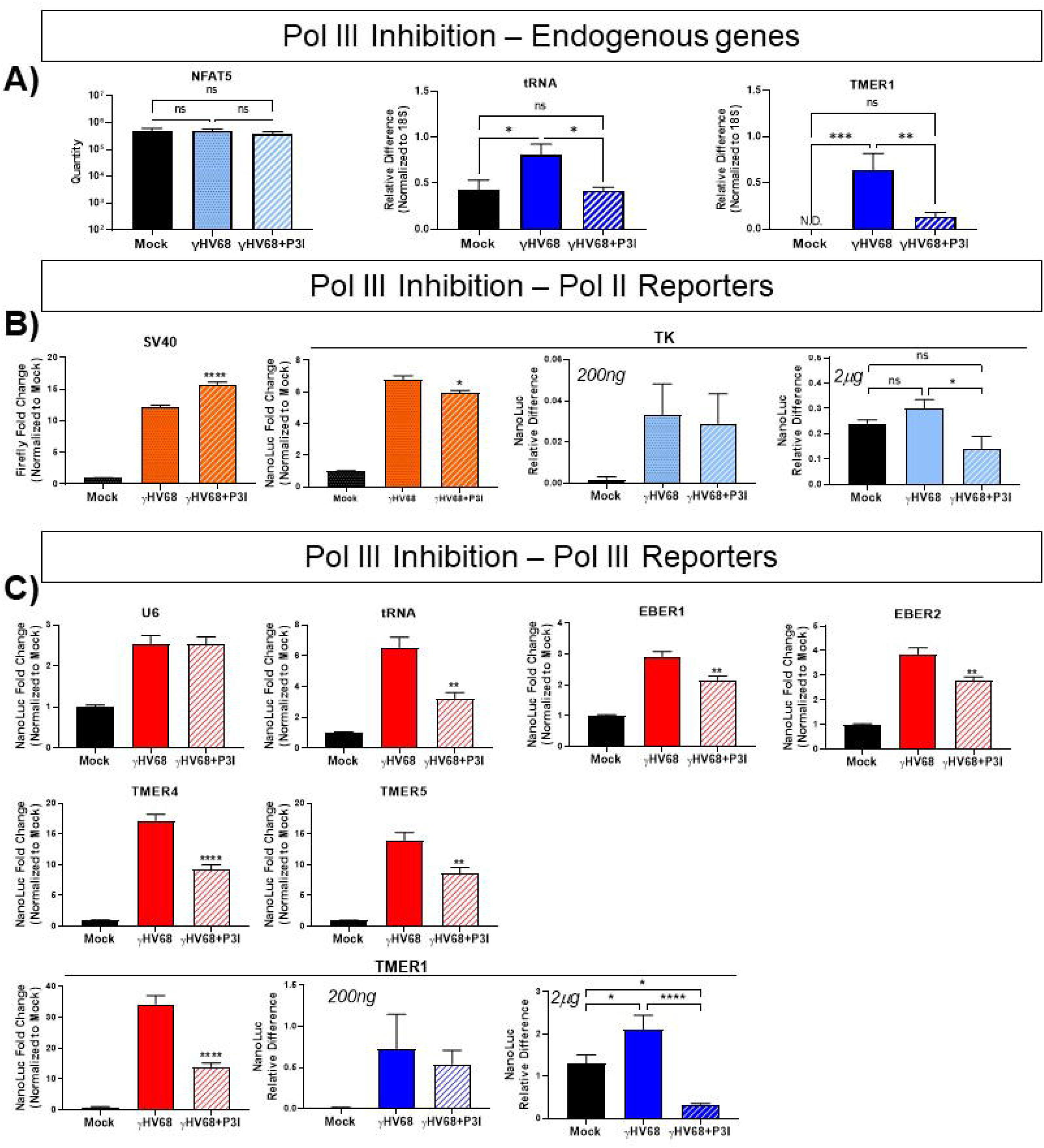
Inhibition of RNA polymerase III decreases luciferase activity and RNA levels from pol III promoter during γHV68 infection. Expression of endogenous control genes and reporter constructs were measured during treatment with a pol III inhibitor. Prior to transfection, samples were treated with an RNA polymerase III inhibitor (CAS 577784-91-9), marked as P3I, or left untreated. Cells were co-transfected with the SV40 promoter driven Firefly luciferase control and NanoLuc reporter constructs with the indicated promoters. Inhibited cells were treated again immediately following infection. Luciferase activity is shown in orange for pol II promoters and red for pol III promoters. RNA expression is shown in light blue for pol II promoters and dark blue for pol III promoters. **A)** Efficacy of pol III inhibition was shown by TaqMan RT-qPCR of a pol II-transcribed gene (NFAT5) and SYBER Green qPCR of pol III-transcribed genes (Human pre-tRNA-Tyr-GTA-1-1 and γHV68 TMER1). Data is from 3 independent experiments with biological duplicates or triplicates per experiment. Luciferase assays were performed as previously described, with cell lysates collected 24 h post-infection. Luciferase activities of the **B)** pol II promoters (orange, SV40-Firefly and TK-NanoLuc) and **C)** pol III promoter-NanoLuc (red) are shown. All luciferase activity changes are analyzed as the fold change in NanoLuc activity normalized to uninfected samples (Infected NanoLuc LU / Mock NanoLuc LU). NanoLuc RNA was also measured from **B)** a pol II promoter (light blue, TK) and **C)** a pol III promoter (dark blue, TMER1). RT-qPCR was performed for NanoLuc RNA as previously described. Cells were treated as described, and transfected with 200 ng or 2 μg of DNA. RNA was collected 24 hpi and analyzed for expression of NanoLuc RNA. Firefly activity from SV40 was measured from two independent experiments with eight samples per experiment. RT-qPCR data for 200 ng are from two independent experiments with biological triplicates, and data for 2 μg are from three independent experiments with biological triplicates or duplicates. Error bars = SEM. Significant differences analyzed by t-test and indicated as asterisks. P-values are indicated as follows: * = P ≤ 0.05, ** = P ≤ 0.01, *** = P≤ 0.001, **** = P ≤ 0.0001.

We also measured the luciferase RNA expressed from the TK (pol II) and TMER1 (pol III) promoters during infection with pol III inhibition. When mimicking the transfection conditions used for the dual luciferase assays with 200 ng transfected DNA, we found no significant difference in the expression of NanoLuc from either the TK or TMER1 promoters in any of the examined conditions (Fig 3B, C, in blue). However, given that there is induction of NanoLuc RNA from the TMER1 promoter during infection (Fig 5), we tested whether increasing the amount of transfected reporter DNA may allow for greater distinction in NanoLuc RNA levels under infection and pol III-inhibition conditions. Thus, we repeated these experiments with 2 μg of total transfected DNA. These experiments showed no difference in NanoLuc expressed from the TK promoter during infection or pol III inhibition as compared to mock. However, there was a significant decrease in NanoLuc expressed from the TMER1 promoter during infection with pol III inhibition compared to both mock (4-fold) and to infection (6-fold). These data suggest that the NanoLuc expression from pol III promoters is dependent on pol III activity.

**Figure 5.**
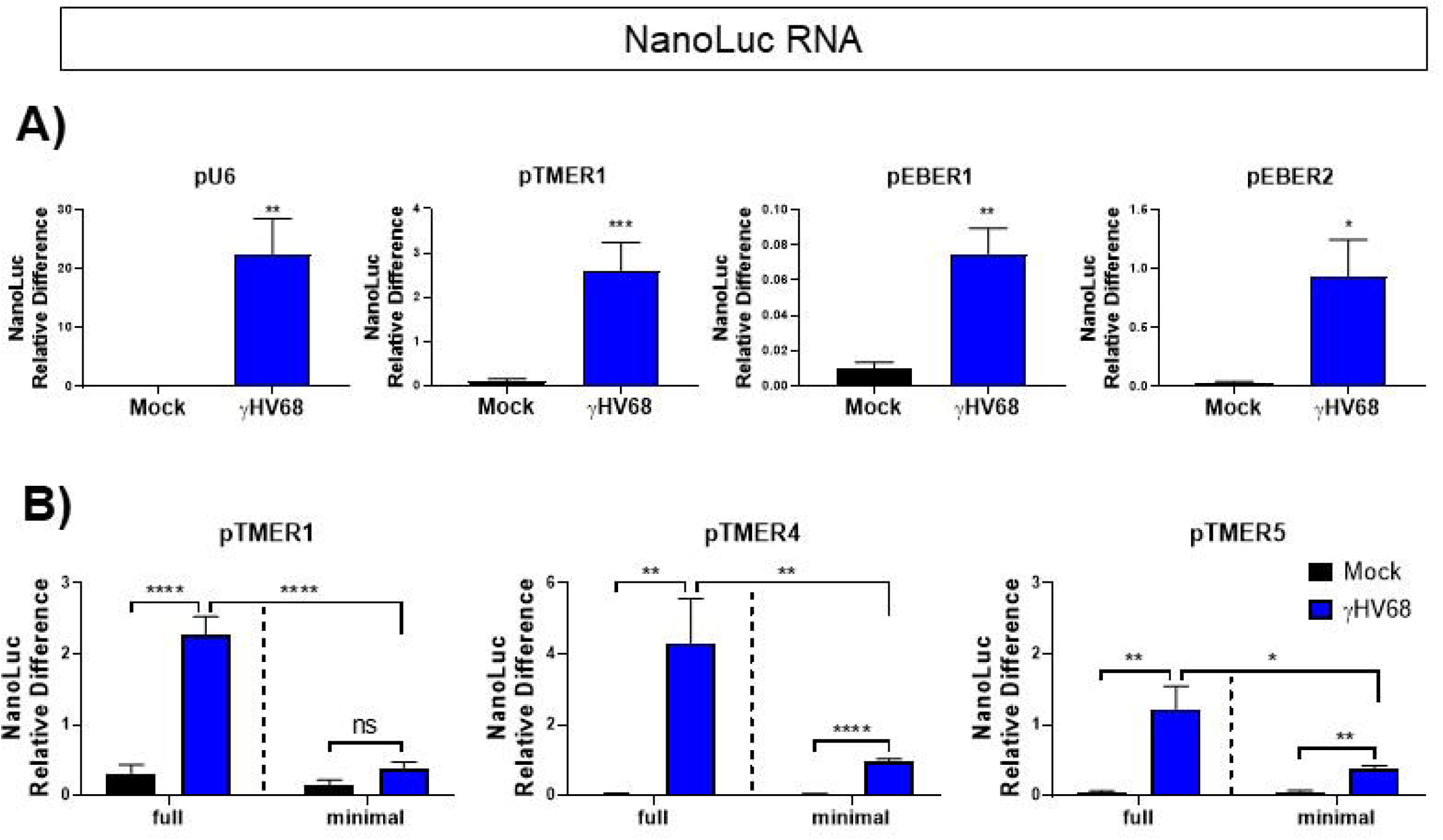
Effect of γHV68 infection on NanoLuc transcript levels. Promoter activity was measured for **(A)** the U6, TMER1, and EBER full promoters, or **(B)** the TMER1, TMER4, and TMER5 minimal vs. full promoters. Cells were treated as previously described and RNA was purified from cells at 16 h post-infection (U6 n = 2, TMER1, n = 5) or 24hpi (EBER1 n = 2, EBER2 n = 2, TMER4 n = 2, and TMER5, n = 2). RNA was converted to cDNA, then LightCycler real-time PCR using Syber Green was performed with primers targeting the NanoLuc gene and a host control gene (18S). The relative difference of NanoLuc was calculated using the Pfaffl method (2001, *Nucleic Acids Research*), where the ratio = (*E*_target_)^ΔCP target (control-sample)^ / (*E*_ref_)^ΔCP ref (control-sample)^. Each experiment (n) included biological triplicates or duplicates. Error bars = SEM. Significant differences analyzed by t-test and indicated as asterisks. P-values are indicated as follows: * = P ≤ 0.05, ** = P ≤ 0.01, *** = P≤ 0.001, **** = P ≤ 0.0001.

Our analyses to this point indicated that the TMER promoters expressed the highest induction in luciferase activity during infection; therefore, we compared the sequences of TMER promoters to identify which features of these promoters could potentially contribute to this strong induction. The initially analyzed TMER promoters contained the TMER promoter, as well as extended sequence around the minimal promoter elements (Fig 4A). Considering that the extra sequence included in these “full” promoters may contribute to infection-induced activity, we created a panel of “minimal” TMER and EBER promoters that contain only the minimal RNA pol III promoter elements, i.e. the sequence beginning from the A box to the end of the B box (Fig 4A). We initially compared the activity of these promoters under basal (no infection) conditions to calculate the average RLUs (NanoLuc LU/Firefly LU) in the absence of virus-induced changes. This analysis showed that U6 was the most active of all pol III promoters under uninfected conditions, followed by the “full” EBER promoters (Fig 4B). In contrast, “minimal” EBER promoters showed a significant reduction in baseline luciferase activity. All TMER promoters (full or minimal) appeared similar to the empty vector, that is, to have virtually no activity under uninfected conditions. We then compared the induction of NanoLuc luciferase activity between the EBER and TMER minimal and full promoters during infection to determine the role of extended sequence on promoter activity. As previously described, these constructs were transfected into HEK 293 cells, then infected with γHV68. The fold change in NanoLuc activity relative to mock-treated samples was compared after 24 h of infection (Fig 4C). When we compared the relative inducibility of EBER “full” versus “minimal” promoters, “minimal” promoters showed greater virus-inducibility. This enhanced inducibility of the “minimal” EBER promoters likely reflects the reduced basal luminescence from these promoters (Fig 4B). Conversely, TMER “minimal” promoters displayed a weaker induction during infection than their “full” counterparts, suggesting the sequence surrounding the TMER minimal promoters drives stronger expression during infection. These results indicate that the sequence surrounding minimal pol III promoter elements impacts both the basal activity and inducibility of these promoters during infection.

**Figure 4.**
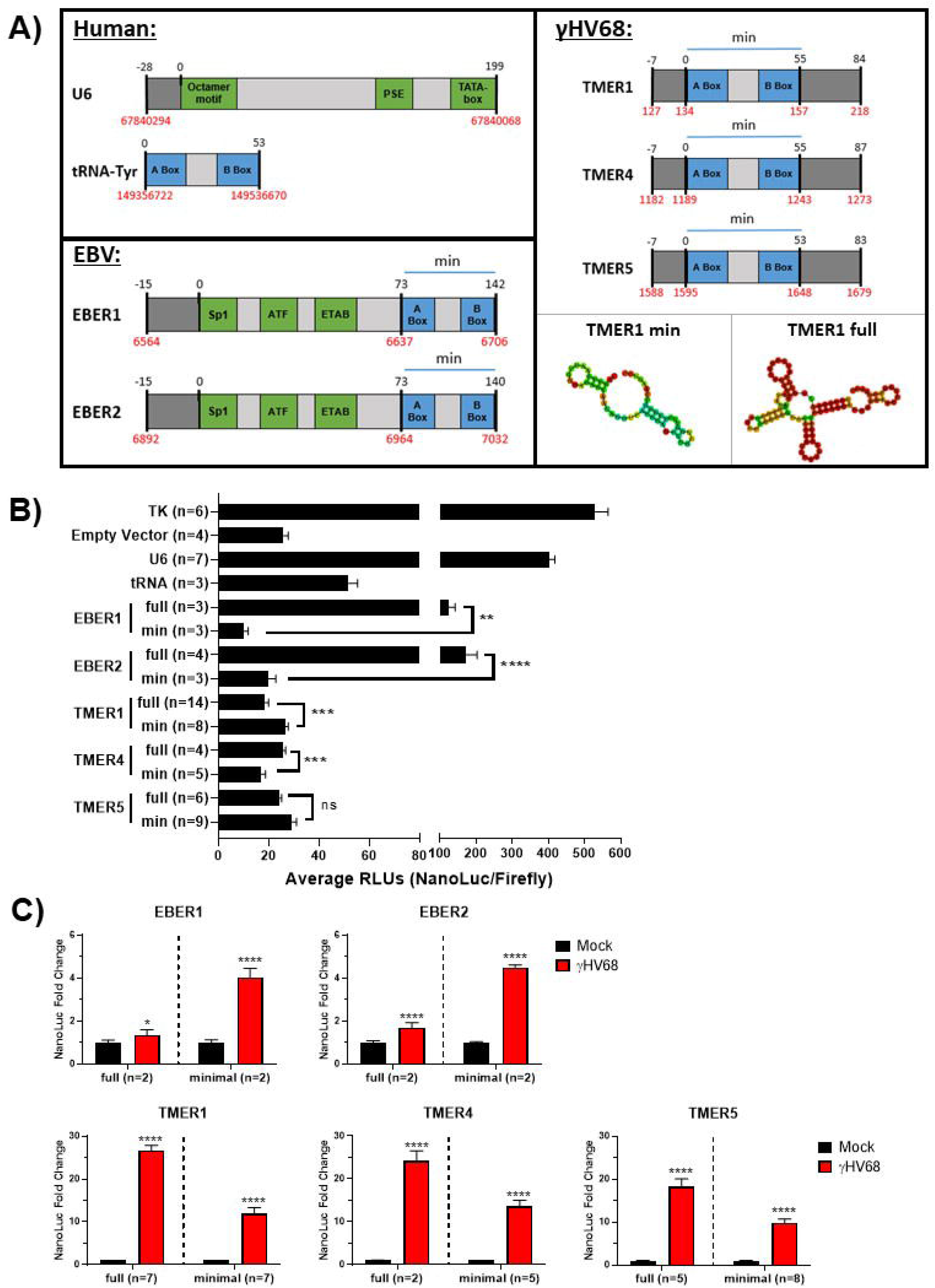
Sequence external to the minimal promoter elements alters promoter activity. **(A)** Several pol III promoters were cloned into the pNLP3 NanoLuc-expressing vector. Promoter schematics are not to scale. The canonical minimal promoter elements for each promoter are shown in blue and notated with a blue line labeled “min”, with the intermediate sequence in light grey. Upstream elements are green, and external/extended sequence is shown in dark grey. Nucleotide positions are shown above each schematic in relation to position “0”, which indicates the start of the minimal promoter sequence. Genomic coordinates are shown below each schematic in red, and refer to the following reference sequences; Human U6 (RNU6-1) = NC_000015.10, Human tRNA-Tyr (TRY-GTA11-1) = NC_000007.14, EBERs in Human Herpesvirus 4/EBV Whole Genome = NC_007605.1, TMERs in MHV68 Whole Genome = NC_001826.2. The coordinates for human U6 snRNA and tRNA are reversed to reflect their orientation as coded on the complement strand. The predicted structure for the minimal and full TMER1 promoter using the RNAfold Webserver are shown. **(B)** Cloned reporter constructs were transfected into HEK293 cells at a 10:1 molar ratio with a Firefly luciferase control expressed by the SV40 promoter. TK-expressed NanoLuc was used as a positive control. Cells were not infected. Following 48 hours of transfection, dual luciferase assays were performed as previously described. Luciferase activity is expressed as the ratio of NanoLuc activity to control Firefly activity (relative luminescence units, RLUs). **(C)** Luciferase assays were performed as previously described, with cell lysates collected 24 h post-infection. All promoter activity changes are analyzed as the fold change in NanoLuc activity normalized to uninfected samples. Each experiment (n) included biological triplicates. Error bars = SEM. Significant differences analyzed by t-test and indicated as asterisks. P-values are indicated as follows: * = P ≤ 0.05, ** = P ≤ 0.01, *** = P≤ 0.001, **** = P ≤ 0.0001.

Luciferase readouts of pol III promoter activity allowed us to uniformly analyze pol III promoter activity. This assay does not directly measure the level of RNAs, however, instead relying on an enzymatic readout of luciferase protein activity. To ensure that γHV68 infection was inducing pol III activity transcriptionally, we used the same NanoLuc constructs to measure promoter activity at the RNA level by performing RT-qPCR for the NanoLuc transcript. Following the same protocol as used for the luciferase assays, HEK 293 cells were transfected with pGL3 and the pNLP3 vector expressed by pol III promoters of interest (as outlined in Fig 1). Cells were then infected with γHV68 and RNA was purified from cells 16 or 24 h post-infection. Primers targeting the NanoLuc gene were used for qPCR following reverse transcription of the RNA. Infection increased the NanoLuc RNA expression from the U6 and TMER1 promoters, with more modest induction from the EBER promoters (Fig 5A). These results indicate that γHV68 infection stimulates pol III-promoter activity from multiple host and viral promoters, measured at both the transcriptional and translational level. To extend these findings, we further measured NanoLuc RNA expression from “minimal” or “full” TMER promoters. These studies demonstrated that γHV68 infection increased NanoLuc RNA from the “minimal” promoter relative to mock infected samples, with further RNA induction from the “full” TMER promoter (Fig 5B). These results strongly suggest that the NanoLuc reporter assay serves as a faithful readout for pol III-dependent transcription, quantified at both the RNA and protein level. These findings also emphasize that sequences outside of the minimal TMER promoters contribute to increased expression during infection.

γHV lytic replication critically depends on viral DNA replication and late gene transcription, processes that are inhibited by phosphonoacetic acid (PAA) (42, 43). We therefore tested the impact of PAA on virus-induced pol III induction. To do this, HEK 293 cells were transfected with the pol III-driven NanoLuc constructs and infected as before, with one set of samples receiving PAA treatment (200μg/mL) after 1 h of viral inoculation. PAA treatment was consistently associated with increase luciferase enzymatic activity, with PAA-treated γHV68-infected cultures characterized by a greater apparent induction of luciferase activity compared to γHV68-infected cultures alone. This PAA-driven enhancement of luciferase activity was observed for multiple pol III promoters, including U6, TMER1, 4 and 5, and EBER 1 and 2 (Fig 6A). To determine if this effect was also observed at the transcriptional level, cells were transfected and infected as before. RNA was isolated 16 h post-infection and RT-qPCR was performed to detect the NanoLuc transcript. Notably, treatment with PAA during γHV68 infection had no impact on the induction of NanoLuc RNA compared to untreated infected cells, indicating that viral DNA replication and late gene synthesis was not required for pol III induction (Fig 6B). Effectiveness of the PAA treatment was confirmed by measuring expression of a viral late gene, gB (Fig 6C). The increase in luciferase activity following PAA treatment, with minimal impact on NanoLuc RNA, strongly suggests that PAA treatment enhanced the translational output from the promoters tested. These data suggest that viral late gene expression plays an additional role in translation that is not seen at the transcriptional level, a phenomenon independent of pol III promoter activity.

**Figure 6.**
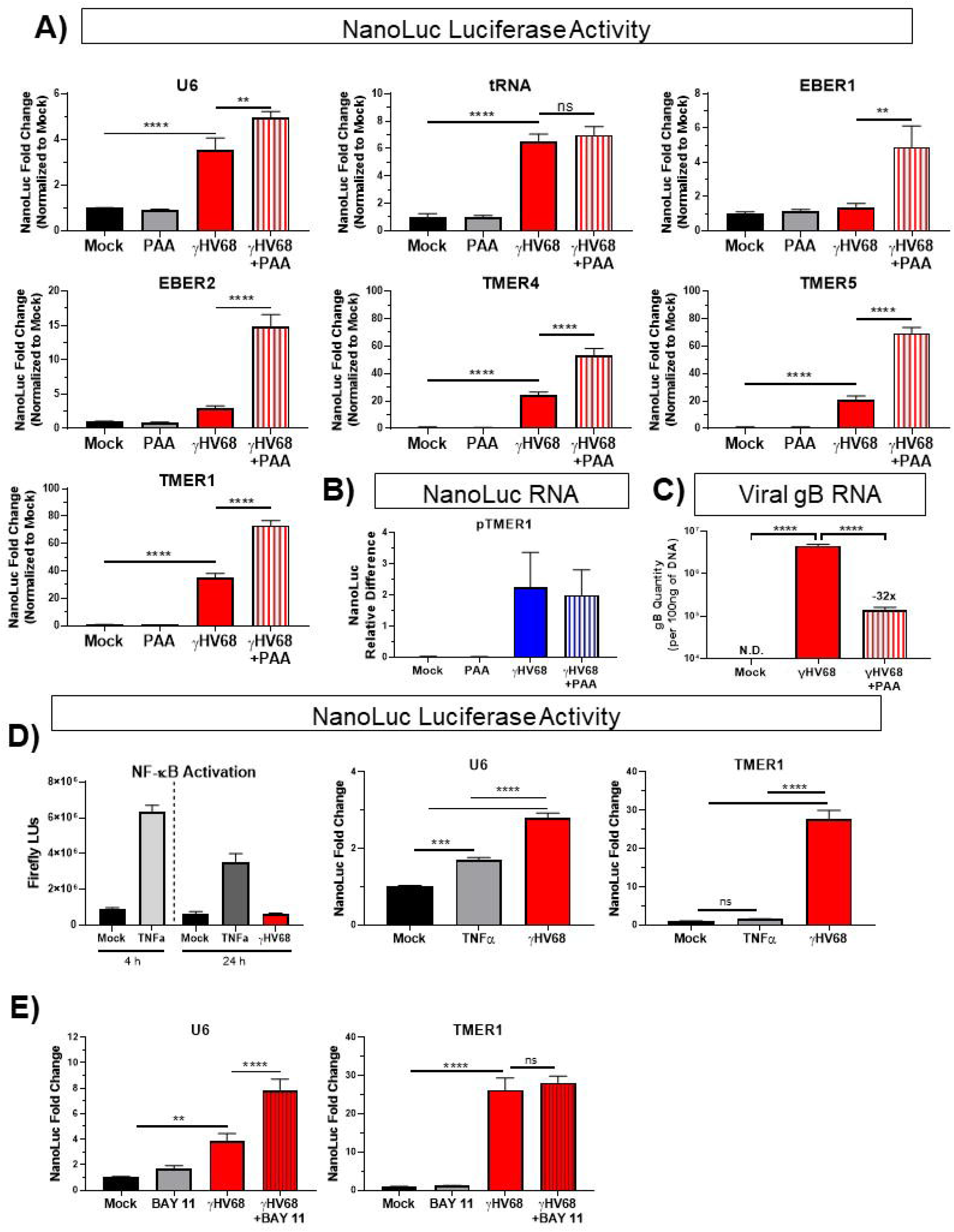
Effect of viral and NF-κB inhibitors and an NF-κB activator on luciferase activity during γHV68 infection. **A)** Promoter activity was measured via luciferase activity for the following promoters during infection, including with concurrent PAA treatment: TMER1 (n = 5), TMER4 (n = 2), TMER5 (n = 2), EBER1 (n = 2), EBER2 (n = 3), U6 (n = 2), and tRNA (n = 1). Each experiment contained biological duplicates or triplicates. Luciferase assays were performed as previously described, with cell lysates collected 24 h post-infection. NanoLuc fold change for infected samples is from previous figures for comparison to PAA-treated samples. All promoter activity changes are analyzed as the fold change in NanoLuc activity normalized to uninfected samples. **B)** Promoter activity was measured via RT-qPCR of the NanoLuc transcript expressed from the TMER1 promoter during infection and concurrent phosphonoacetic acid (PAA) treatment. RT-qPCR was performed as previously described. Data is shown as NanoLuc relative difference to a host gene (18S). N = 2 with biological triplicates. **C)** Using the same RNA from experiments shown in panel B, RT-qPCR was performed for a viral late gene (gB) to confirm the efficacy of PAA treatment, which resulted in an approximately 32-fold decrease in gB compared to WT infected samples. gB was not detected (N.D.) in mock samples. **D)** NF-κB activation with 50 ng/μL TNFα (NF-κB reporter n = 1 for each time point, U6 n = 2, TMER1 n = 3) or **E)** NF-κB inhibition by 10 μM BAY 11-7082 (U6 n = 3, TMER1 n = 4). NF-κB activation is shown in Firefly luminescence units, while NanoLuc activity from the U6 or TMER1 promoters is expressed as fold change over mock. Cells included in the TNFα experiments were transfected for 12 h, while all others were transfected for 24 h as previously described. Each experiment (n) contains biological triplicates. Error bars = SEM. Significant differences analyzed by t-test (two conditions) or one-way ANOVA (three conditions) and indicated as asterisks. P-values are indicated as follows: * = P ≤ 0.05, ** = P ≤ 0.01, *** = P≤ 0.001, **** = P ≤ 0.0001.

Given the reported relationship between the NF-κB pathway and the expression of pol III-dependent transcripts (30), we analyzed the effect of NF-κB activation or inhibition on the activity of the U6 and TMER1 promoters via luciferase activity. First, we measured induction of an NF-κB reporter plasmid following treatment with either TNFα, a known inducer of the NF-κB pathway, or following γHV68 infection. Whereas TNFα induced NF-κB reporter activity at 4 and 24 hours post-treatment, γHV68 infection had no measurable impact on expression from the NF-κB reporter (Fig 6D). Next, we analyzed the impact of NF-κB manipulation on pol III promoter activity. Treating cells with TNFα modestly increased U6 promoter activity, albeit to a lesser extent than γHV68 infection, while TMER1 promoter activity was not affected by TNFα treatment (Fig 6D). Inhibition of NF-κB with the BAY 11-7082 (BAY 11) compound increased U6-expressed luciferase activity in virus infected conditions, yet had no significant impact on TMER1 promoter activity after infection (Fig 6E). This indicates a potential role of NF-κB in inhibiting pol III promoter activity during infection; however, this effect is only observed in the case of a gene-external (i.e. type 3) promoter. Ultimately, these data do not support a significant role of the NF-κB pathway in the observed induction of pol III promoter activity after γHV68 infection in these culture conditions.

The NanoLuc-expressing constructs allowed us to measure promoter activity using a reporter system including a shared readout that minimizes confounding factors of RNA sequence/structure/stability, and our analysis of pol III promoter activity suggested a general induction during infection. However, a previous report suggested that only a subset of host pol III-transcribed genes – the SINE RNAs – are increased during lytic γHV68 infection of NIH3T3 cells and *in vivo* infections of C57BL/6 mice (33). While our reporter studies rely on cell systems that can be easily transfected with reporter DNA, we wanted to quantify the impact of γHV68 infection on the abundance of endogenous murine and viral pol III RNAs, yet avoid challenges in PCR amplification, unique primer/probe designs, and bulk analysis. In order to accomplish this, we made use of the PrimeFlow assay system, a sensitive and robust fluorescent in situ hybridization assay, combined with multiparameter flow cytometry. This system quantifies steady-state RNA expression at the single-cell level using direct probe hybridization to endogenous RNAs, with sensitivity provided by amplification based on probe stacking rather than PCR and primers. For this analysis, murine fibroblast cells (3T12) were mock-treated, infected with wild-type (WT) γHV68, or infected with an EBER-knock in (EBER-KI.γHV68) recombinant γHV68 that lacks the TMERs and instead contains insertion of the EBV EBERs into the TMER locus. EBER-KI.γHV68 was competent for viral replication (manuscript in progress). 3T12 cells were infected with an MOI of 1 and harvested at 16 h post-infection, conditions that result in a mixed population of virally infected and uninfected cells. Cells were then queried with fluorescent probes to detect viral (the γHV68 TMERs or the EBV EBERs) and host ncRNAs (U6 snRNA or 4.5 rRNA, the murine equivalent of human 5S rRNA). This analysis allowed a comparison of host ncRNA expression as a function of viral ncRNA expression, comparing cells with or without virus ncRNA expression. Gating schemes for doublet exclusion and probe specificity are shown in Figure 7. Notably, probes for the TMERs and EBERs were specific for their intended targets, with no detectable TMER or EBER signal in mock-infected samples, TMER+ cells only present in WT γHV68 infected samples, and EBER+ cells only present in EBER-KI.γHV68 infected samples (Fig 8E). We next compared the relative expression of the U6 (Fig 8A-B) and 4.5S ncRNAs (Fig 8C-D) between cells with active viral RNA expression versus those cells that did not express these viral ncRNAs. Analysis of the geometric mean fluorescence intensity (gMFI) for the U6 snRNA and 4.5 rRNA probes revealed increased expression of U6 and 4.5 rRNA ncRNAs in virally infected cells (i.e. TMER+ or EBER+) compared to uninfected cells (TMER- or EBER-) (Fig 8F-G). These data demonstrate an increase in host pol III-transcribed ncRNAs in γHV68-infected cells and emphasize the benefit of single-cell analysis to quantify endogenous ncRNA expression during virus infection.

**Figure 7.**
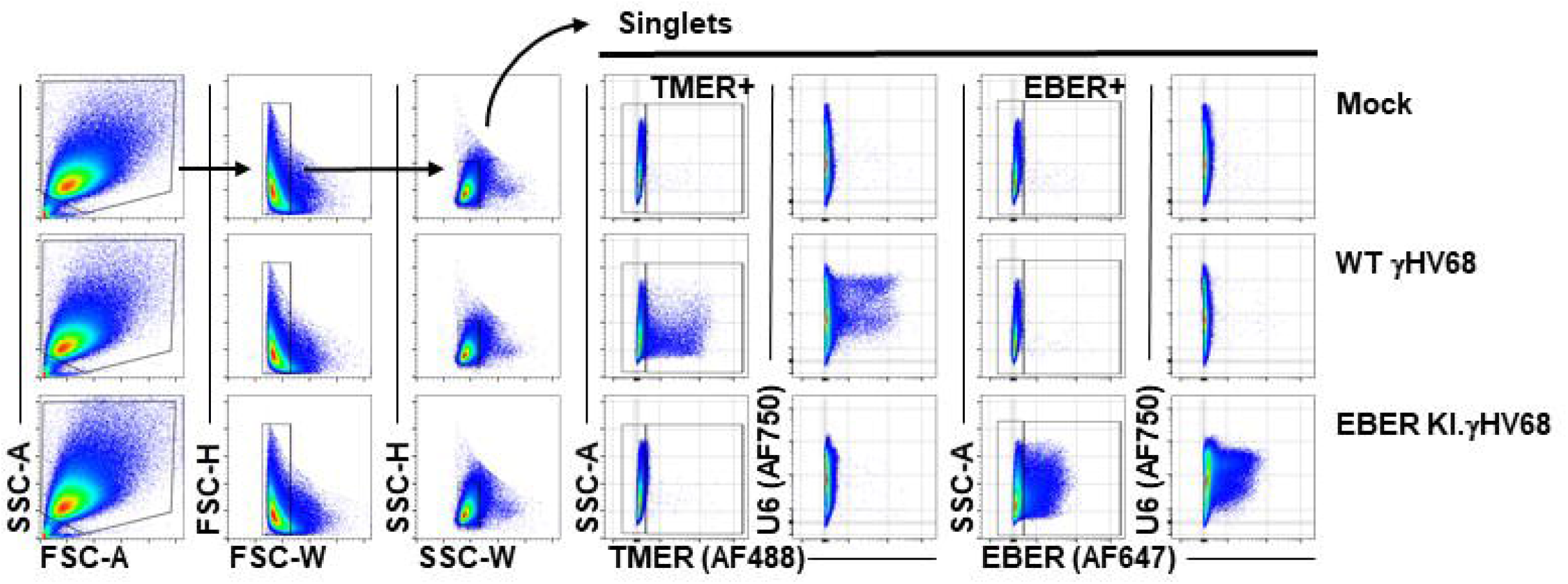
Viral ncRNA expression serves as an indicator of virus infection. Murine 3T12 fibroblasts were infected with mock, WT γHV68, or an EBER knock-in (EBER-KI) γHV68 at an MOI=1, harvested at 16 hpi, and subjected to the PrimeFlow RNA Assay and flow cytometric analysis. Singlets were identified using a sequential gating strategy as shown, with representative examples from mock (top), WT γHV68 (middle) or EBER-KI.γHV68 infected samples. Singlet populations were then analyzed for TMERs (Type 4/AF488), EBERs (Type1/AF647), and U6 snRNA or 4.5S RNA (Type 6/AF750 for both, each in different probe mixes).

**Figure 8.**
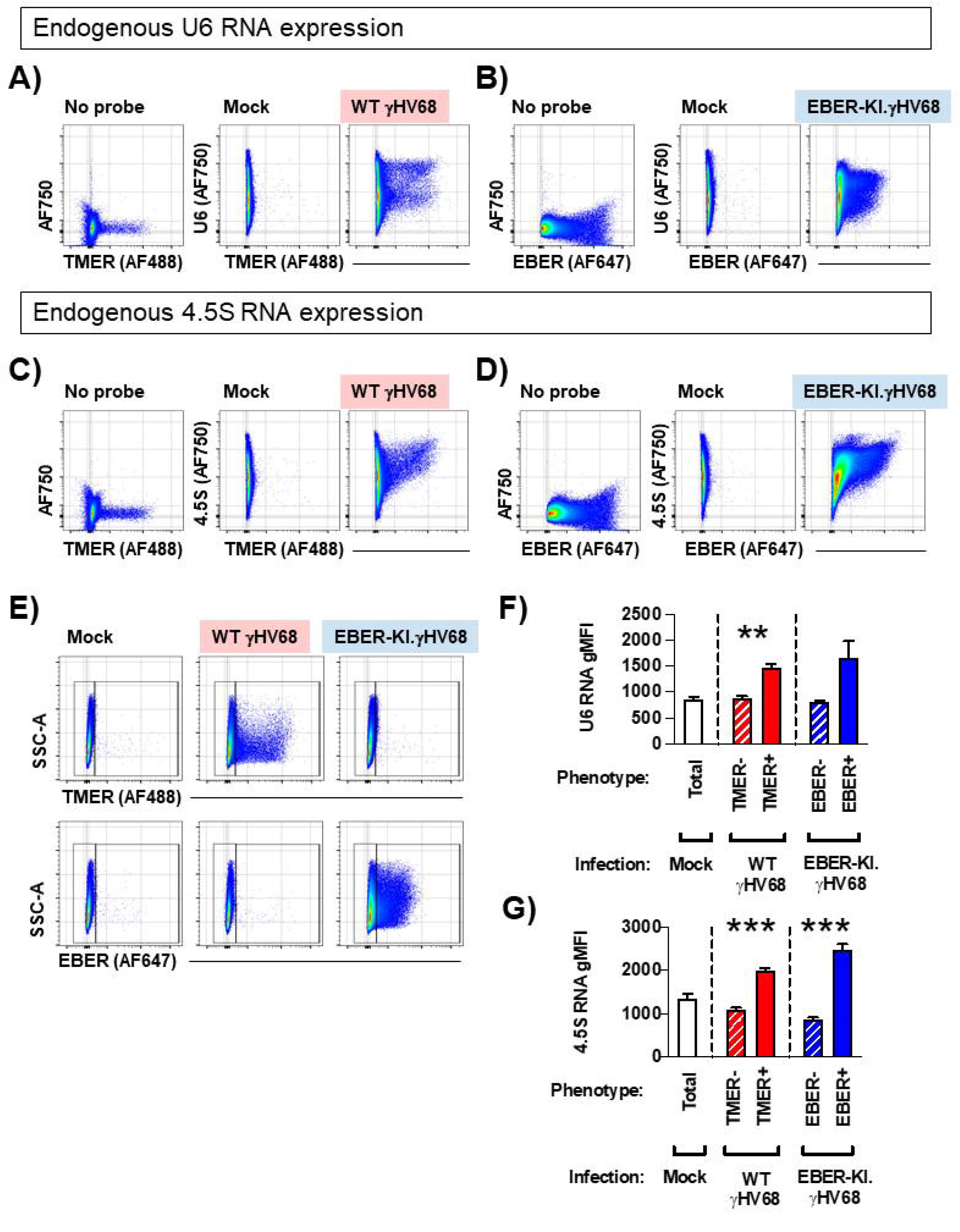
Endogenous expression of murine U6 snRNA and 4.5S rRNA during γHV68 infection. Murine fibroblast cells (3T12) were infected with mock, WT γHV68, or an EBER knock-in (EBER-KI) γHV68 at an MOI=1, harvested at 16 hpi and subjected to the PrimeFlow RNA Assay and flow cytometric analysis. **(A**,**B)** Analysis of U6 snRNA expression comparing background fluorescence (”No probe”, left) versus U6 probe hybridization in Mock (middle), or WT γHV68 or EBER-KI.γHV68 infected cells (right). **(C**,**D)** Analysis of 4.5S rRNA expression comparing background fluorescence (”No probe”, left) versus 4.5S rRNA probe hybridization in Mock (middle), or WT γHV68 or EBER-KI.γHV68 infected cells (right). **(E)** Identification of virally-infected cells that express either the TMERs or the EBERs, following infection with mock, WT γHV68 or EBER-KI.γHV68. Gates define cells based on positive or negative expression as indicated, with these populations used for subsequent quantitation in panels F and G **(F**,**G)** The geometric mean fluorescence intensity (gMFI) was calculated for **(F)** U6 snRNA or **(G)** 4.5S rRNA in mock infected, or virus infected samples, comparing cells that differed in expression of either the TMERs or the EBERs in WT or EBER-KI infections respectively. Data depict flow cytometric analysis of singlets, defined by sequential discrimination using SSC and FSC parameters as shown in Figure 7. Data are from 3 biological replicates, with statistical significance assessed using an unpaired t-test. Statistical significance as follows: **, p<0.01 and ***, p<0.001.

The above analysis established a correlation between expression of viral and host ncRNAs at the single-cell level, but raised the potential that detection of endogenous viral ncRNAs could bias our analysis to cells with higher pol III activity independent of viral effects. To address this concern, we repeated the analysis with another, pol III-independent read-out of infection, detection of the pol II-dependent γHV68 ORF18 RNA. Cells were treated as before, with the modification that they were infected at an MOI of 5 rather than 1 so that differences between infected and non-infected cells were exaggerated. To simplify the RNA probe panel, and to fortify previous data, we focused on the U6 snRNA as a host measure of pol III activity. Infected cells were defined as ORF18 RNA positive, as well as TMER (Fig 9A) or EBER (Fig 9B) positive for WT and EBER-KI γHV68 infection, respectively. This analysis identified several populations that could be distinguished by different expression levels (negative, mid or high expression) for either ORF18 or the viral ncRNAs (Fig 9C-D). Within each ORF18-defined population, we calculated the gMFI for either the TMERs/EBERs (Fig 9E) or U6 snRNA (Fig 9F). ORF18 RNA^high^ cells had the highest expression of the TMERs, in WT infected samples, and the EBERs, in EBER-KI infected samples compared to ORF18 RNA^mid^ or ORF18 negative cells (Fig 9E). Notably, virally-infected ORF18 RNA^high^ cells from either WT or EBER-KI γHV68 infection were also characterized by increased U6 expression relative to uninfected, ORF18 negative cells in the same cultures (Fig. 9F). Consistent with our previous observations, U6 RNA gMFI was highest in cells with highest expression of either the TMERs or EBERs (Fig 9G). These data demonstrate that virally infected cultures are characterized by variation in ncRNA expression between individual cells, and that cells with abundant virus transcription are further characterized by increased host and viral ncRNA expression.

**Figure 9.**
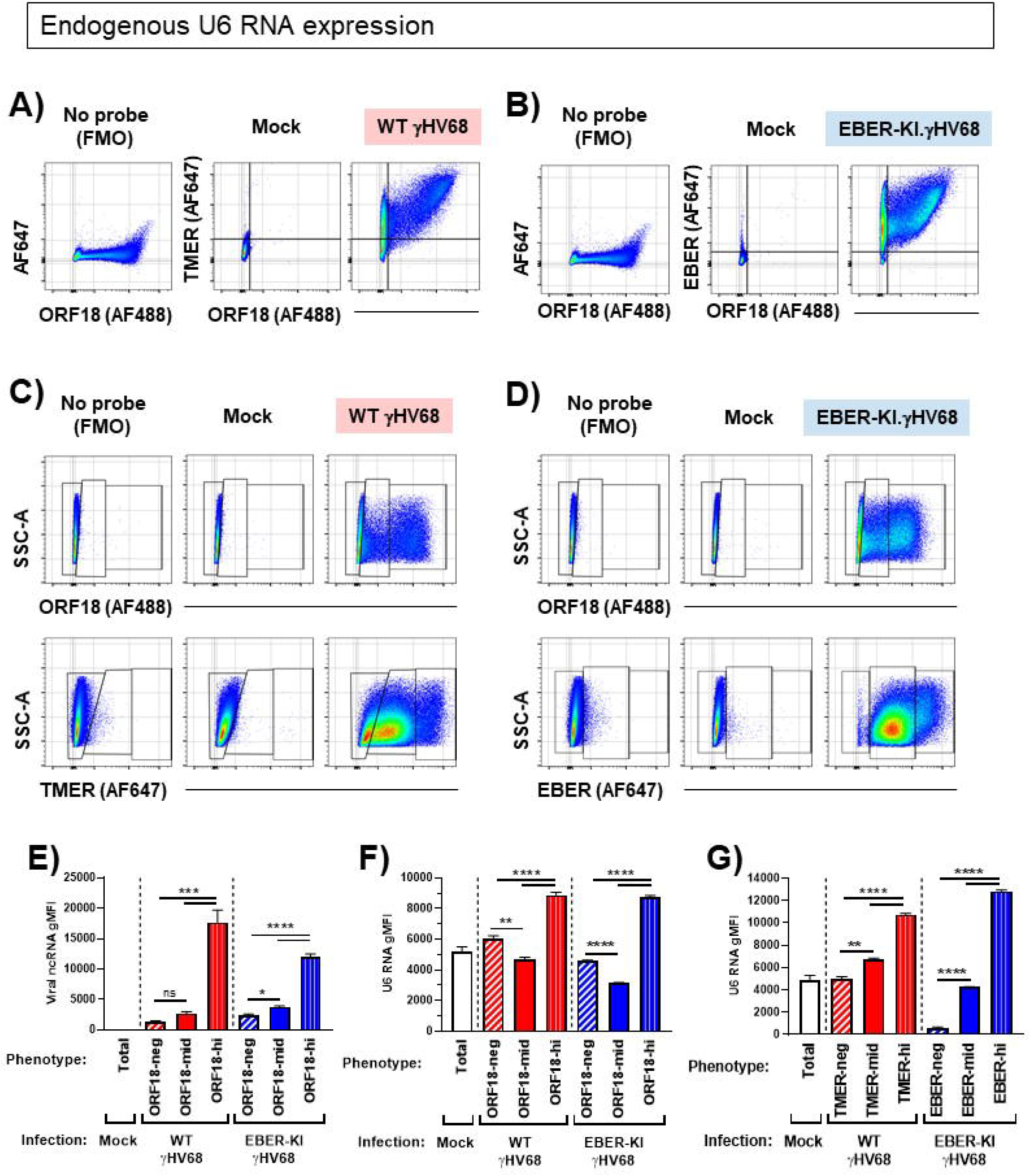
Endogenous expression of murine U6 snRNA during γHV68 infection. Murine fibroblast cells (3T12) were infected with mock, WT γHV68, or an EBER knock-in (EBER-KI) γHV68 at an MOI=5, harvested at 16 hpi and subjected to the PrimeFlow RNA Assay and flow cytometric analysis. **(A**,**B)** Analysis of viral gene expression indicating viral ncRNA (TMER or EBER) expression and/or ORF18 expression comparing background fluorescence (“No probe”, left) versus TMER or EBER hybridization in Mock (middle), or WT γHV68 or EBER-KI.γHV68 infected cells (right). **(C**,**D)** Analysis of viral gene expression indicating ORF18 expression (top) or viral ncRNA (TMER or EBER, bottom) expression comparing background fluorescence (“No probe”, left) versus probe hybridization in Mock (middle), or C) WT γHV68 or D) EBER-KI.γHV68 infected cells (right). These plots indicate the gating used for further analysis in E-F, and gating is duplicating from Figure 9. **(E)** The geometric mean fluorescence intensity (gMFI) was calculated for viral ncRNA (TMERs or EBERs) in mock infected or virus infected samples, comparing cells that differed in expression of ORF18. **(F**,**G)** The gMFI was calculated for U6 snRNA in mock or virus infected samples, comparing cells that differed in expression of either **(F)** ORF18 or **(G)** TMERs or EBERs in WT or EBER-KI infections. Data depict flow cytometric analysis of singlets, defined by sequential discrimination using SSC and FSC parameters as exemplified in Figure 7. Data are representative of two experiments, with 3 biological replicates in each. Statistical significance assessed using a one-way ANOVA with multiple comparisons. Statistical significance as follows: * = P ≤ 0.05, ** = P ≤ 0.01, *** = P≤ 0.001, **** = P ≤ 0.0001.

## DISCUSSION

While γHV infection is known to alter expression of host genes, and viral ncRNAs are integral for pathogenesis, the transcriptional regulation of these ncRNAs has remained unclear. Here, we propose that different pol III promoter types allow for distinct means for transcriptional regulation during infection. This study focused on the impact of γHV68 lytic replication on pol III promoter activity. To measure pol III activity, we used three highly-sensitive methods, two of which were based on a reporter gene and one method based on measuring authentic RNAs. Though these methods each have different advantages and disadvantages, they all indicate that lytic infection drives a general upregulation of promoter activity across multiple host and viral pol III-dependent transcripts, as shown by upregulation of both luciferase activity and luciferase RNA expression, as well as increased expression of endogenous transcripts as measured by flow cytometric analysis of pol III RNAs. Our studies further reveal distinct effects of specific promoters and promoter features. These findings emphasize the utility of the modified NanoLuc luciferase system to analyze pol III promoter activity, and provide clear evidence for pol III promoters with large differences in basal and inducible promoter activity. They further emphasize the capacity of γHV lytic infection to modify pol III-dependent transcriptional machinery in infected cells, a process that likely facilitates productive virus replication (33). At this time, it remains unknown whether pol III machinery or transcription is altered during γHV68 latent infection or reactivation from latency.

Our use of a luciferase reporter to measure the activity of pol III promoters allowed us to directly compare the functional activity of multiple pol III promoters in a high-throughput assay while minimizing potential differences that may arise due to ncRNA sequence, structure, or stability. While these assays used NanoLuc RNA and protein as a standard platform for analysis, an important caveat of this analysis is that many pol III promoters are gene-internal. Based on this, the resulting transcript is comprised of a hybrid of at least the minimal pol III promoter elements (i.e. the A and B box) directly fused upstream of the NanoLuc RNA, creating a hybrid RNA that is not 100% identical between constructs. Though pol III promoters conventionally drive expression of non-coding RNAs, there is clear precedent that pol III can transcribe translation-competent RNA (44, 45), and luciferase reporters have been used for high throughput and unbiased analysis of pol III promoter activity (38, 46). Inspired by these studies, we cloned several host and viral ncRNA promoters into a NanoLuc luciferase reporter to measure their activity during lytic γHV68 infection. We chose the NanoLuc luciferase as our reporter due to the small size and high activity, which is consistent with pol III transcriptional capacity and measurement of genes that may be expressed at a low level. To further enhance the robustness of this reporter, we identified and removed a pol III termination sequence within the NanoLuc gene which approximately doubled luciferase reporter activity. Though pol III transcription should theoretically be terminated in the original NanoLuc reporter, pol III read-through of termination signals has been reported (47). In total, use of the modified NanoLuc reporter construct afforded a sensitive and robust readout for assessing pol III promoter activity.

Through use of this pol III reporter assay, we found that γHV68 infection increased promoter activity across a range of host and viral pol III promoters as measured by both luciferase activity and reporter RNA abundance. Pol III transcription from these promoters was confirmed through treatment with a pol III specific inhibitor (CAS 577784-91-9). These results demonstrate that virus induction of pol III promoters is dependent on RNA polymerase III, either through direct or indirect means. Interestingly, the consequence/magnitude of induction elicited by infection varied between promoters. For example, the U6 promoter conveyed high basal activity, with infection resulting in a modest induction of U6 promoter activity. Conversely, the TMER promoters exhibited extremely low basal activity in mock-infected conditions, with dramatic induction after γHV68 infection. The inducibility of the TMERs was further enhanced by accessory sequences outside of the minimal A and B box elements. One explanation for this enhanced induction is that these extended sequences may contain additional transcriptional elements that are integral to the promoter itself. While formally possible, it is notable that accessory sequences across the TMERs are not conserved (6, 22, 23). An alternate explanation for the enhanced activity of the full TMER promoters is that inclusion of the extended sequence includes the full tRNA-like structure of the TMER genes (Fig 4A). It is interesting to speculate that this tRNA-like structure could either lend greater stability to transcripts, or protect the transcripts from degradation by host exonucleases or the viral endonuclease, muSOX (48). Although pol III transcription is frequently associated with the transcription of housekeeping ncRNAs, there is clear precedent that pol III can also participate in inducible gene expression (e.g. γHV68-induced expression of SINEs) (33, 34). Whether the TMERs have conserved regulatory mechanisms with host inducible ncRNAs is currently unknown, however, the TMERs share promoter similarity to the SINEs (type 2, gene internal).

Our studies demonstrated that infection increased NanoLuc expression at both the RNA and protein activity level, indicating that virus infection increased pol III promoter activity and not some secondary measurement. While we saw broad induction of luciferase activity across pol II and pol III promoters, induction of activity from pol III promoters was inhibited by a pol III-inhibitor, consistent with a model that these promoters are recruiting pol III to transcribe translation-competent RNA. Notably, induction of the reporter RNA as measured by RT-qPCR was specific to a pol III promoter (TMER1), and no significant induction was seen from a pol II promoter (TK, Fig 3B). This suggests possible caveats in the induction we measure from luciferase activity vs. luciferase RNA. A difference in reporter protein activity vs. transcript abundance was also seen when we inhibit viral late gene expression. Unexpectedly, inhibition of viral late gene expression with PAA resulted in increased luciferase activity from nearly all of the promoters examined – this phenomenon was only seen at the level of NanoLuc protein activity, not at the level of NanoLuc RNA. The ability of PAA to enhance NanoLuc protein activity, with no commensurate change in RNA expression, suggests that viral late genes may have a potential role in tempering translation. Together with the general induction of luciferase activity seen during infection, these data suggest that viral induction of gene expression may be occurring at both the transcriptional and translational level. Infection may induce expression of pol III-derived reporter transcripts (RNA) and also increase general translation (luciferase activity), leading to a compound effect when we measure reporter protein activity. Manipulation of host translational machinery by the herpesviruses is a common strategy that is required for optimal virus replication and the production of virus progeny (49).

While experiments with the NanoLuc reporter constructs indicated a general induction in the activity of the observed pol III promoters, the levels of induction varied by promoter type. Interestingly, promoters with upstream elements (U6, EBER1, and EBER2) displayed the highest level of basal activity in mock conditions and the lowest level of virus-mediated induction of luciferase activity, while gene-internal type 2 promoters had the lowest basal activity and highest induction (Fig 2). This is likely not due to promoters with gene external elements reaching the limit of detection of the luciferase assay, as demonstrated by the increased induction seen with PAA treatment (Fig 6). Notably, the luciferase activity expressed from the U6 promoter was not significantly impacted by pol III inhibition, while all other pol III promoters showed decreased activity (Fig 3C). Among the promoters with external elements, the U6 promoter has the highest induction of NanoLuc expression when measured at the RNA level and the EBERs had a relatively low induction, with the gene-internal TMERs moderately induced (Fig 5). The inhibitors used in this project also appear to have promoter-specific affects, with NF-κB modulation altering the output from the type 1 U6 promoters but not the type 2 TMER promoter (Fig 6). Though all transcripts are transcribed by pol III, these data indicate varying induction of the different promoters, with varying abilities for the resulting transcripts to be translated into functional proteins. As evidenced by the U6 promoter, the external elements of type 1 pol III promoters seem to drive unique responses compared to other pol III promoters and these unique characteristics must be kept in mind when analyzing different promoter types.

Due to the possibility that viral infection has unique effects on transfected plasmids, we complemented our reporter studies with a highly sensitive analysis of endogenous and viral gene expression using the PrimeFlow assay to quantify endogenous RNAs by flow cytometry. Notably, this assay allowed us to identify gene expression differences within individual cells that may be lost in bulk analysis. Furthermore, the use of a branching fluorescent probe targeting the RNA of interest allows magnification of low signals without requiring replication of the gene of interest with potentially inadequate/inefficient primers. These data were consistent with our previous findings, indicating that viral infection lead to the induction of both viral (TMERs, EBERs) and host (U6 snRNA, 4.5 rRNA) pol III-derived transcripts. An alternate explanation for these findings is that cells with inherently higher pol III activity are more susceptible to viral infection, resulting in the observation of both high viral pol II gene expression (ORF18) coupled with high expression of pol III-derived ncRNAs. As this assay measures steady-state RNA levels that are influenced by RNA production and decay, changes in measured RNA levels may be influenced by factors such as transcriptional induction, RNA stability, and/or degradation.

Previous reports show that γHV infection can have diverse effects on pol III transcription, ranging from a general induction of pol III machinery (e.g. in the context of EBV and EBNA1 (32, 50)), to the selective induction of pol III-transcribed RNAs (e.g. induction of specific host vault RNAs in EBV latently infected cells (30, 31), and the host SINE RNAs in γHV68-infected cells (33)). One challenge in interpreting these different findings is that these studies have been done in different states of infection (latent versus lytic), in different cell types, and in different states of cellular transformation. In many cases, the mechanistic insights gained from these studies could only be gained through the use of *in vitro* studies. In keeping with this, we anticipate that the γHV68 system will afford unique insights in how the γHVs regulate pol III-dependent ncRNA expression, allowing the analysis of primary infection coupled with technologies to measure promoter activity and endogenous ncRNA expression. For example, our single-cell analysis of pol III-derived transcripts – U6 snRNA and 4.5 rRNA – supported that γHV68-infection not only increases pol III-dependent promoter activity (as observed in experiments with reporter constructs), but also increases the endogenous expression of these transcripts. Future strategies to improve high throughput direct comparison of promoters could include the use of reporter based constructs such as those described in this study along with other reporter-based readouts, such as recently reported fluorescent RNA aptamers that do not rely on translation (51). How host and viral ncRNAs are regulated as a function of cell type and virus stage of infection remains an important unanswered question.

In total, our studies revealed a γHV68-dependent induction in the activity of host and viral pol III promoters. This induction was seen in the expression of a reporter gene, as well as in the endogenous expression of pol III-dependent transcripts. Though previous reports have focused on the virus-mediated upregulation of specific host ncRNAs, these experiments suggest a broader effect of lytic γHV68 infection on pol III activity. This suggests that γHV68 modulation of the host transcriptional landscape goes beyond mRNA regulation, and that pol III-dependent transcripts are likely to play a wider role in γHV68 pathogenesis than previously appreciated.

## METHODS

### Viruses and tissue culture

All viruses were derived from a BAC-derived WT γHV68 (52). For some experiments, the TMER total knock-out (TMER-TKO) virus was used; this virus was generated as previously described (25). Viruses were propagated and titered as previously described (25). EBER-KI virus contains EBERs 1 and 2 in the TMER-TKO virus backbone. Its generation and characterization are described in a manuscript in preparation.

Human endothelial kidney (HEK 293) and murine fibroblast (3T12) cells were cultured in Dulbecco’s modified Eagle medium (DMEM; Life Technologies) supplemented with 5% fetal bovine serum (FBS; Atlanta Biologicals), 2 mM L-glutamine, 10 U/ml penicillin, and 10 μg/ml streptomycin sulfate (complete DMEM). Cells were cultured at 37°C with 5% CO_2_.

### Mutagenesis of pNL1.1 to create the pNLP3 NanoLuc luciferase reporter

The promoterless NanoLuc luciferase reporter vector pNL1.1[*Nluc*] was obtained from Promega, and primers were designed to introduce silent mutations to remove the pol III termination signal in the NanoLuc coding sequence; these primers are listed in Table 2. Mutagenesis PCR was performed with the following cycles: (i) 95° for 30s, (ii) 12 cycles of 95°C for 30s, 55°C for 1 min, 68°C for 3 m. The resulting DNA was digested with DpnI (New England Biolabs Inc) and transformed into XL1-Blue super-competent cells (Agilent). Bacterial colonies were sequenced to confirm the correct mutations. The resulting plasmid was named “pNLP3” to indicate that it is a <u>N</u>ano<u>L</u>uc plasmid optimized for <u>p</u>ol <u>III</u>.

**Table 1.**
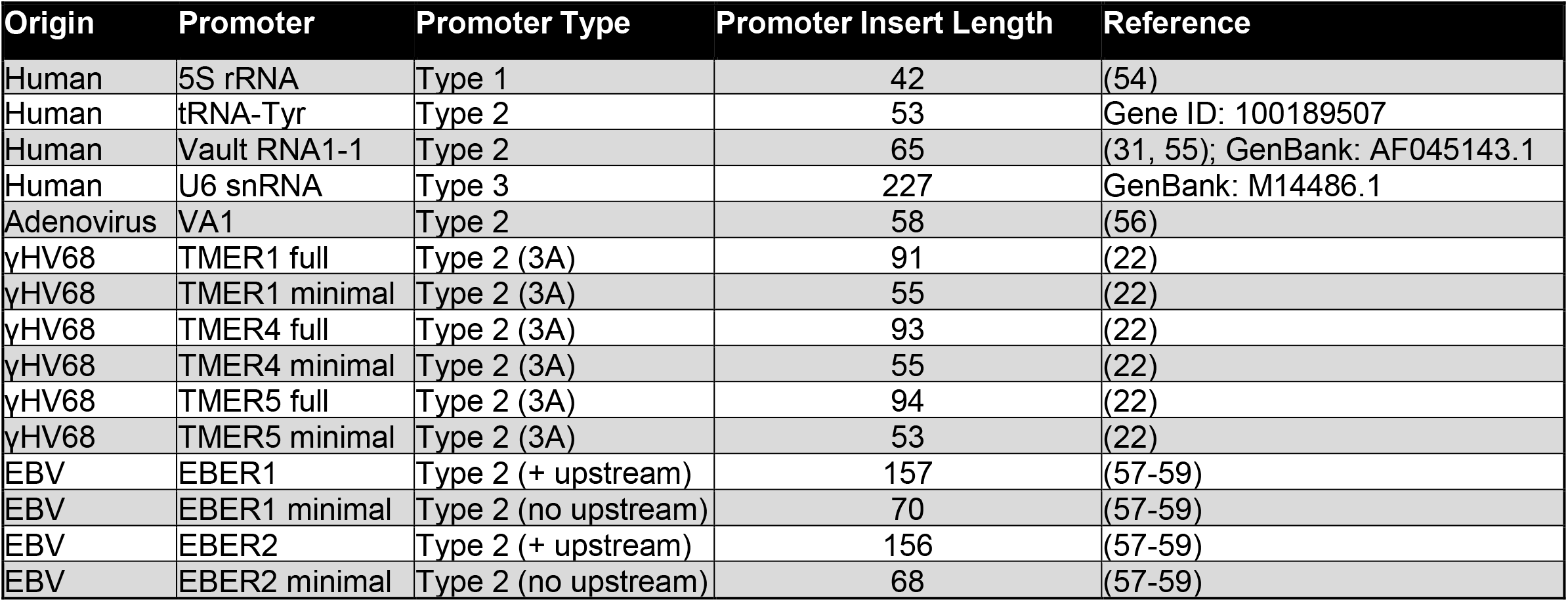
Promoters analyzed in this study. Pol III promoters were selected from several organisms and cloned into a luciferase reporter to analyze promoter activity during infection. Parenthetical additions specify promoter features (“3A” = triplicated A box; for EBERs, “+ upstream” = includes upstream elements, “no upstream” = excludes upstream elements).

| Origin | Promoter | Promoter Type | Promoter Insert Length | Reference |
| --- | --- | --- | --- | --- |
| Human | 5S rRNA | Type 1 | 42 | (54) |
| Human | tRNA-Tyr | Type 2 | 53 | Gene ID: 100189507 |
| Human | Vault RNA1-1 | Type 2 | 65 | (31, 55); GenBank: AF045143.1 |
| Human | U6 snRNA | Type 3 | 227 | GenBank: M14486.1 |
| Adenovirus | VA1 | Type 2 | 58 | (56) |
| γHV68 | TMER1 full | Type 2 (3A) | 91 | (22) |
| γHV68 | TMER1 minimal | Type 2 (3A) | 55 | (22) |
| γHV68 | TMER4 full | Type 2 (3A) | 93 | (22) |
| γHV68 | TMER4 minimal | Type 2 (3A) | 55 | (22) |
| γHV68 | TMER5 full | Type 2 (3A) | 94 | (22) |
| γHV68 | TMER5 minimal | Type 2 (3A) | 53 | (22) |
| EBV | EBER1 | Type 2 (+ upstream) | 157 | (57-59) |
| EBV | EBER1 minimal | Type 2 (no upstream) | 70 | (57-59) |
| EBV | EBER2 | Type 2 (+ upstream) | 156 | (57-59) |
| EBV | EBER2 minimal | Type 2 (no upstream) | 68 | (57-59) |

**Table 2.**
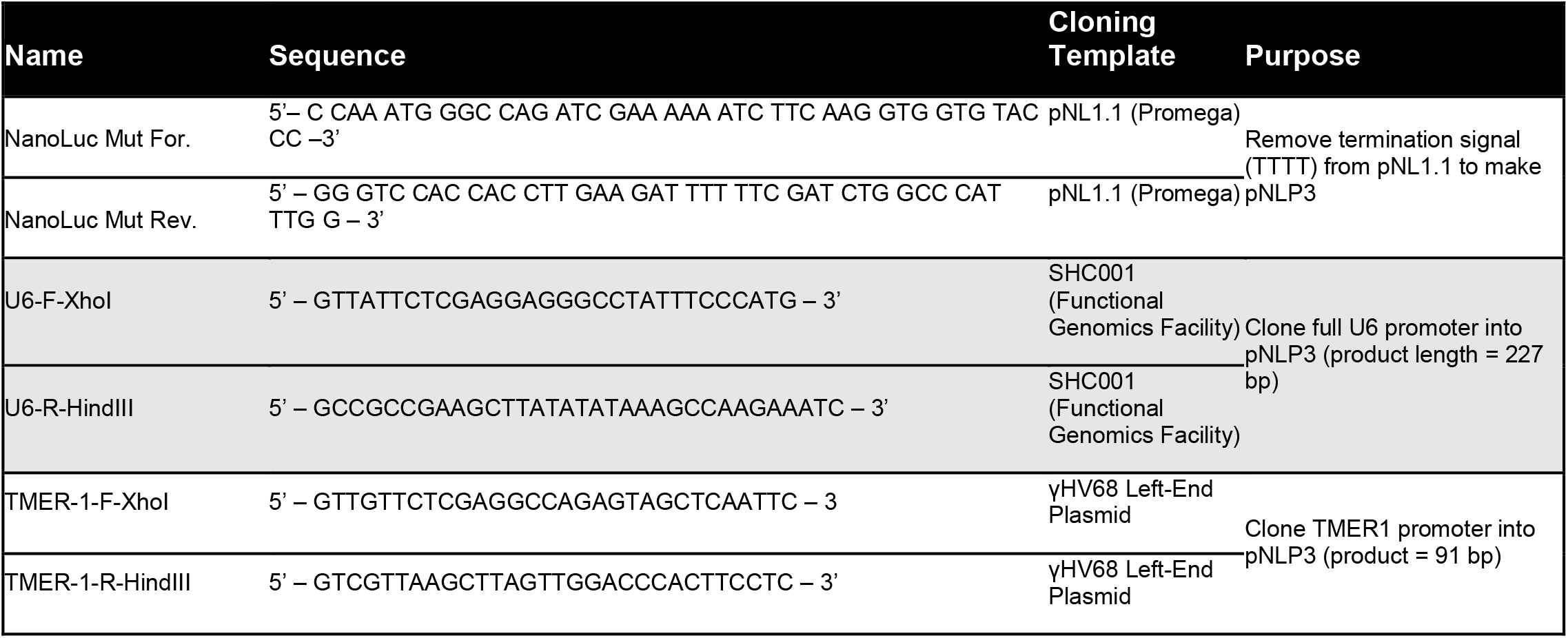
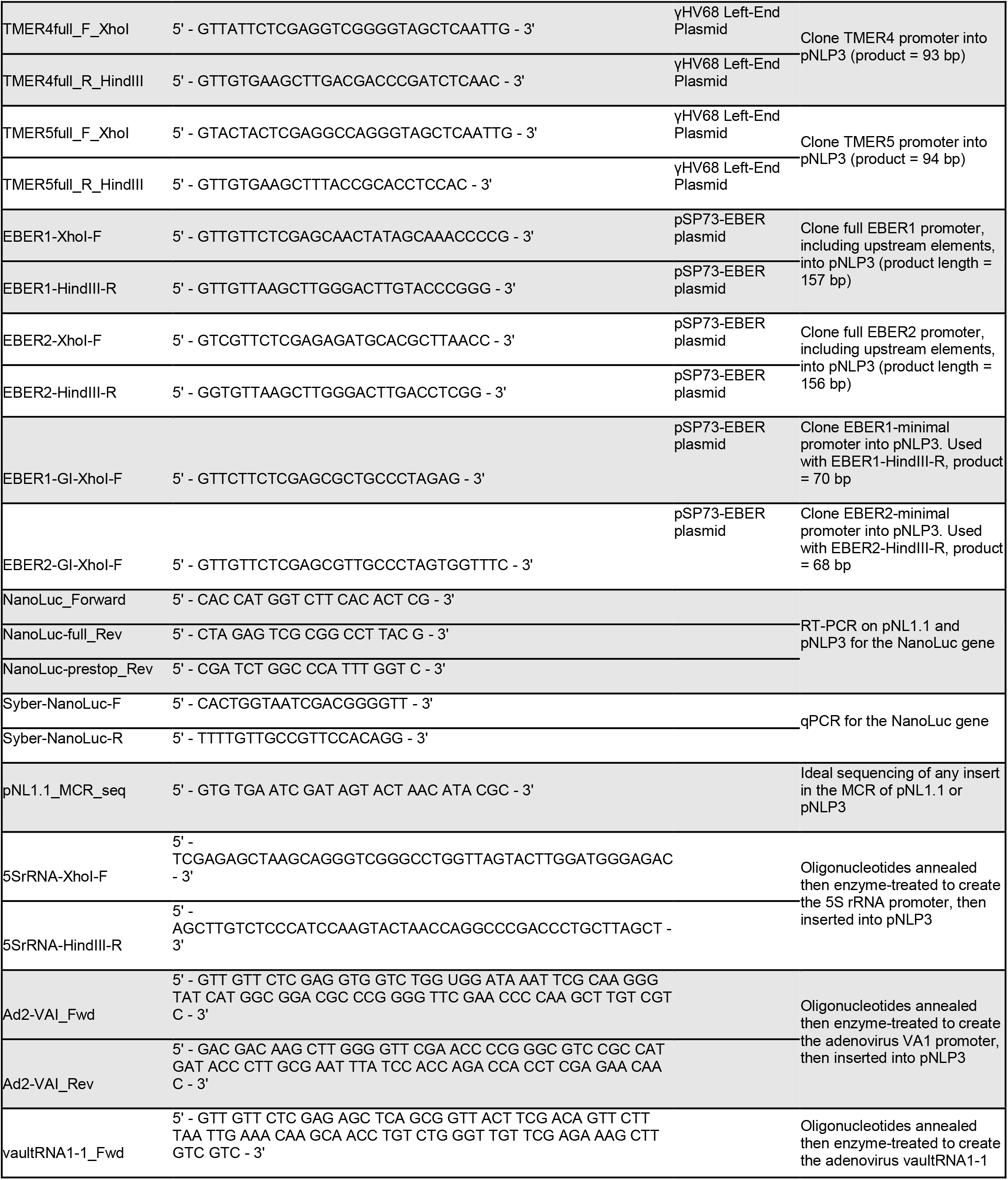
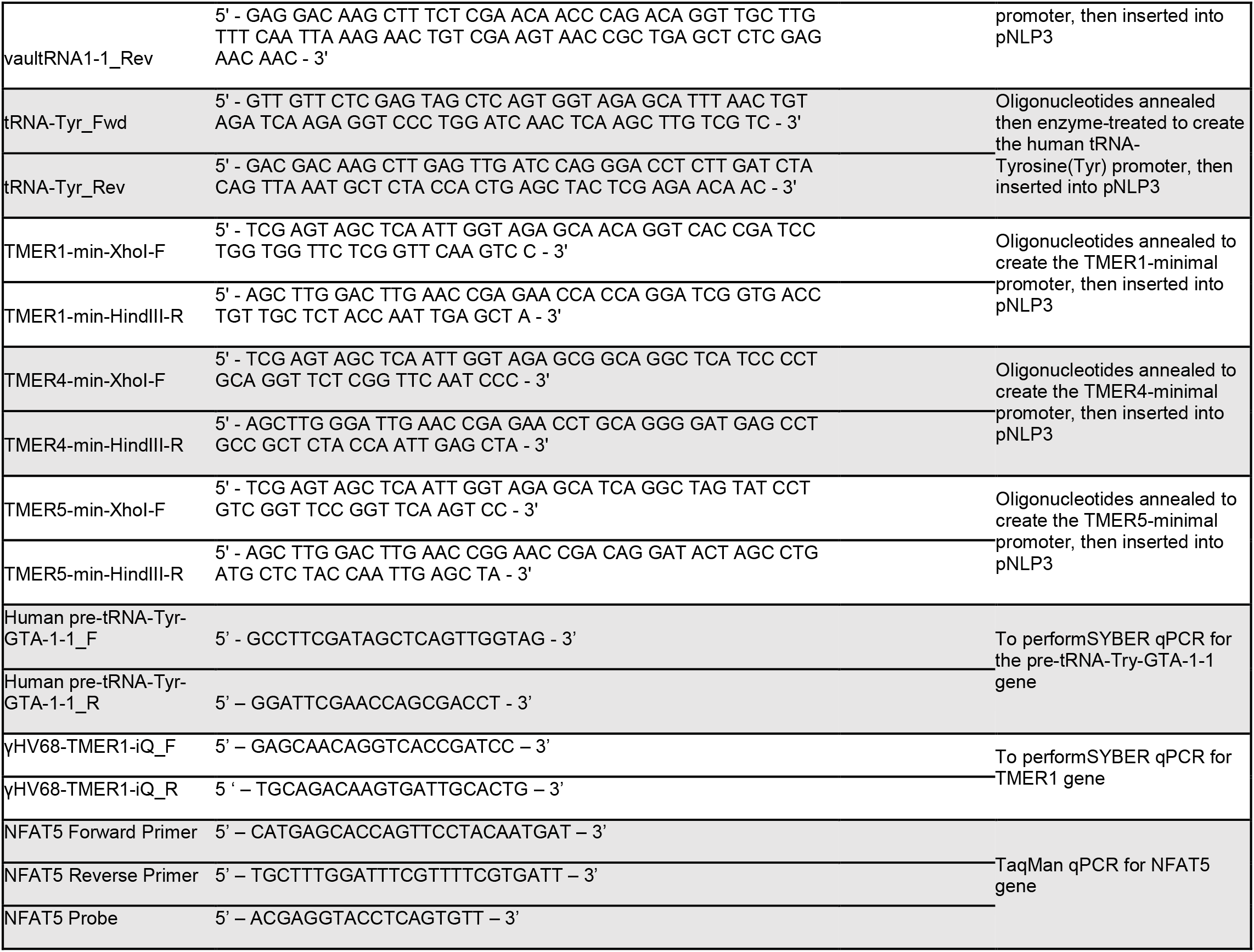
Sequences of primers and oligonucleotides used in this study. Includes sequences for primers, oligonucleotides, and probes used for cloning reporter constructs, sequencing, RT-PCR, and qPCR.

### Generating a pol III promoter-driven NanoLuc reporter panel

All promoters were generated to include XhoI and HindIII overhang sequences on the 5’ and 3’ ends respectively. Several promoters were constructed using ligated oligonucleotides. All sequences for primers and oligonucleotides used in this work are shown in Table 2. PCR-amplified promoters and pNLP3 were digested with XhoI and HindIII, then promoters were ligated into pNLP3 using T4 DNA ligase (New England BioLabs Inc). Ligated constructs were transformed into One Shot electro- or chemically competent TOP10 *E. coli* cells (Thermo Fisher Scientific, #C404052 or #C404010), which were then plated at several dilutions on LB agar containing ampicillin. Resulting colonies were expanded in LB broth with ampicillin and plasmid was isolated using the QIAprep Spin Miniprep Kit (Qiagen). All constructs were confirmed by sequencing.

### Transfecting cells

For transfections, HEK 293 cells were cultured in 5% FBS DMEM without penicillin or streptomycin for approximately 24 hours. Transfection solutions contained Opti-MEM (Thermo Fisher Scientific), NanoLuc plasmid (pNLP3 with inserted pol III promoters), and the Firefly control plasmid (pGL3-Control; Promega). After plasmids were added to the Opti-MEM, solutions were incubated with X-tremeGENE HP DNA Transfection Reagent (Sigma-Aldrich) for at least 15 minutes at room temperature. Transfections were performed in several plate formats, depending on the downstream use; transfection solutions were added dropwise to the appropriate wells (10 μL of solution for 96-well plate, 100 μL of solution in 12-well plates, 200 μL of solution in 6-well plates). For all transfections, the molar ratio of NanoLuc plasmid to Firefly plasmid was kept at 10:1; the total amount of DNA transfected per well was adjusted depending on the plate size (approximately 10 ng for 96-well, 100 ng for 12-well, and 200 ng for 6-well). Some experiments involved transfecting 2 μg of total plasmid per well in a 6-well format. Cells were incubated with transfection solution for 24 hours prior to downstream applications, unless otherwise stated.

To analyze how promoter activity and pol III transcription was affected by γHV68 infection, transfected HEK 293 (for NanoLuc experiments) or 3T12 (for RNA flow) cells were infected with WT, TMER-TKO, or EBER-KI γHV68 at a multiplicity of infection (MOI) of 1 or 5 plaque forming units per cell. Cells were cultured for approximately 24 h prior to infection. Cell counts were determined by treating with 0.05% Trypsin-EDTA (Life Tech, #25300-054) to remove and cells. These cells were mixed with Trypan Blue dye (Bio-Rad, #145-0021) to obtain a live cell count using the TC20 Automated Cell Counter (Bio-Rad). Virus stocks were mixed with 5% complete DMEM, then added to cells and incubated with virus for 1 hour. Viral inoculum was then removed and replaced with complete 5% DMEM. For samples treated with phosphonoacetic acid (PAA, 200 μg/mL, Sigma-Aldrich #284270), inoculum was removed and replaced with 5% complete DMEM containing the drug. Inhibition of RNA polymerase III was achieved by treating cells with 40 μM of CAS 577784-91-9 (Sigma, #557404-M) immediately prior to transfection and again following viral inoculation.

### Dual luciferase assays

Following transfection and infection, HEK 293 cell lysates were collected for analysis of luciferase activity. All luciferase assays were performed using the Nano-Glo® Dual-Luciferase® Reporter Assay System (Promega). For experiments performed in the 12-well format, supernatant was removed and cells were scraped, collected into 1.5mL tubes, then pelleted. Cell pellets were resuspended in 250 μL of Passive Lysis Buffer, per the manufacturer’s protocol. 80 μL of the cell suspensions were used for assays. For luciferase assays performed in the 96-well format, 80 μL of ONE-Glo^™^ EX Reagent was added directly to the cells and supernatant, as recommended by the manufacturer. Samples were incubated at room temperature while on a shaker for three minutes. This solution was then transferred from a transparent 96-well culture plate to a white-wall luminescence plate prior to reading Firefly luciferase activity on the Tecan Infinite® 200 PRO plate reader. Then, 80 μL of the NanoDLR^™^ Stop & Glo® Reagent was added to the solution and incubated at room temperature on a shaker for at least 10 minutes. Samples were read on the Infinite® 200 PRO again for the NanoLuc luminescence measurement.

### RT-PCR and RT-qPCR

RNA was isolated from transfected HEK 293 cells with TRIzol® Reagent (Thermo Fisher Scientific, #15596026) and DNase-treated with TURBO^™^ DNase (Invitrogen, #AM2238) following the manufacturer’s protocols. RNA amplification and removal of DNA was confirmed by RT-PCR or PCR amplification of the NanoLuc gene and a control host gene, 18S. RT-PCR was performed using the OneStep RT-PCR Kit (Qiagen, #210212), with the following conditions: (i) 50°C for 30 m, (ii) 95°C for 15 m, (iii) 40 cycles of 94°C for 30 s, 52°C for 30 s, and 72°C for 30 s, (iv) 72°C for 10 m, (v) hold at 4°C. PCR was performed using Taq DNA polymerase (Qiagen, #201205) with the following conditions: (i) 95°C for 5 min, (ii) 40 cycles of 94°C for 30 s, 52°C for 30 s, 72°C for 30 s, (iii) 72°C for 10 m, (iv) hold at 4°C. RNA samples that showed no product following PCR amplification were deemed DNA-free, and converted to cDNA using SuperScript III Reverse Transcriptase (Invitrogen, #18080093) following the manufacturer’s protocol. 100 ng of the cDNA was then used for qPCR analysis of the NanoLuc, human pre-tRNA-Tyr-GTA-1-1, γHV68 TMER1, and human 18S genes using the iQ™ SYBER® Green Supermix (Bio-Rad, #1708880) with the following conditions: i) 95°C for 3 m, ii) 40 cycles of 95°C for 15 s, 60°C or 61°C for 1 m, iii) 95°C for 15 s, 60°C for 1 m, 95°C for 15 s. Amplification of NanoLuc, pre-tRNA, or TMER1 was normalized to 18S expression to calculate the relative difference of target gene expression using the Pfaffl method: 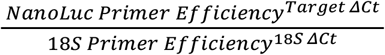 (53). Single product for each target was confirmed by melt curves and gel electrophoresis of product following qPCR.

NFAT5 expression was measured using primer-probe qPCR of 100 ng cDNA with the VIC probe and Black Hole Quencher (BHQ). Conditions for the qPCR were: i) 95°C for 10 m, ii) 40 cycles of 95°C for 10 s, 60°C for 30 s, 72°C for 1 s, iii) 40°C for 30 s. Quantity of NFAT5 was determined using a plasmid-based standard curve.

### PrimeFlow RNA assay

The PrimeFlow RNA assay kit (Thermo Fisher Scientific, # 88-18005-210) was used to analyze expression of non-coding RNAs in mock and γHV68-infected 3T12 cells. Cells were infected at an MOI of 1 or 5 for 16 h, then processed following the manufacturer’s protocol. Probes used were: TMERs (Type 4/AF488 or Type 1/AF647), EBERs (Type 1/AF647), ORF18 (Type 4/AF488), and U6 snRNA or 4.5 rRNA (Type 6/AF750), with compatible probe labels depending on the experiment. Samples were collected on an LSR II Flow Cytometer (BD Biosciences), and included single stain and “full minus one” controls. Flow cytometry data were analyzed using FlowJo software (version 10.6.1), with compensation based on single stained beads and cells. Compensated flow cytometry data were subsequently analyzed for singlet events based on doublet discrimination as exemplified in Figure 7. Distinctions of negative and positive populations were based on control samples as shown in Figures 8 and 9.

### Software and statistical analysis

Statistical analysis and graphing were done in GraphPad Prism (Version 8.0d). Statistical significance was tested by unpaired t test (comparing two conditions), one-way ANOVA (comparing three or more conditions), or two-way ANOVA (comparing grouped data) and subjected to multiple corrections tests using recommended settings in Prism. All flow cytometry data were analyzed in FlowJo (version 10.6.1) with flow cytometry data shown as pseudo-color dot plots.

## ACKNOWLEDGEMENTS

This work was supported by the NIH grants R01AI121300 to LFvD and R21 AI134084 to LFvD and ETC, T32 AI052066 to ANK and CO RNA Biosciences summer internship support to AM.

We thank the members of the Clambey and van Dyk lab for helpful discussions, members of the Colorado RNA Bioscience Initiative for their insights, the Colorado ClinImmune core for flow cytometry services, and the Molecular Biology Core Facility at Anschutz Medical Campus for sequencing services. We also thank the Functional Genomics Facility at University of Colorado and the lab of Dr. Joan Steitz lab for their generous gifts of the pLKO.1-puro plasmid (used to clone the U6 promoter) and the pSP73-EBER plasmid (used to clone the EBER promoters), respectively.

